# Loss of Endothelial TDP-43 Leads to Blood Brain Barrier Defects in Mouse Models of Amyotrophic Lateral Sclerosis and Frontotemporal Dementia

**DOI:** 10.1101/2023.12.13.571184

**Authors:** Ashok Cheemala, Amy L. Kimble, Jordan D. Tyburski, Nathan K. Leclair, Aamir R. Zuberi, Melissa Murphy, Evan R. Jellison, Bo Reese, Xiangyou Hu, Cathleen M. Lutz, Riqiang Yan, Patrick A. Murphy

## Abstract

Loss of nuclear TDP-43 occurs in a wide range of neurodegenerative diseases, and specific mutations in the *TARDBP* gene that encodes the protein are linked to familial Frontal Temporal Lobar Dementia (FTD), and Amyotrophic Lateral Sclerosis (ALS). Although the focus has been on neuronal cell dysfunction caused by TDP-43 variants, *TARDBP* mRNA transcripts are expressed at similar levels in brain endothelial cells (ECs). Since increased permeability across the blood brain barrier (BBB) precedes cognitive decline, we postulated that altered functions of TDP-43 in ECs contributes to BBB dysfunction in neurodegenerative disease. To test this hypothesis, we examined EC function and BBB properties in mice with either knock-in mutations found in ALS/FTLD patients (*TARDBP^G348C^* and *GRN^R493X^*) or EC-specific deletion of TDP-43 throughout the endothelium (*Cdh5(PAC)CreERT2; Tardbp^ff^*) or restricted to brain endothelium (*Slco1c1(BAC)CreERT2; Tardbp^ff^*). We found that *TARDBP^G348C^*mice exhibited increased permeability to 3kDa Texas Red dextran and NHS-biotin, relative to their littermate controls, which could be recapitulated in cultured brain ECs from these mice. Nuclear levels of TDP-43 were reduced*in vitro* and *in vivo* in ECs from *TARDBP^G348C^*mice. This coincided with a reduction in junctional proteins VE-cadherin, claudin-5 and ZO-1 in isolated ECs, supporting a cell autonomous effect on barrier function through a loss of nuclear TDP-43. We further examined two models of *Tardbp* deletion in ECs, and found that the loss of TDP-43 throughout the endothelium led to systemic endothelial activation and permeability. Deletion specifically within the brain endothelium acutely increased BBB permeability, and eventually led to hallmarks of FTD, including fibrin deposition, microglial and astrocyte activation, and behavioral defects. Together, these data show that TDP-43 dysfunction specifically within brain ECs would contribute to the BBB defects observed early in the progression of ALS/FTLD.

## Introduction

Loss of nuclear TDP-43 is a common feature in a wide range of neurodegenerative diseases (1). These include Alzheimer’s disease, Limbic-predominant Age-related TDP-43 Encephalopathy (LATE), and ALS/FTD. Most notably, in FTD, a significant subset of the disease is characterized by the aggregation of uniquinityled-TDP-43 in the cytosol and nuclear exclusion. The identification of familial FTD mutations in TDP-43 which exacerbate this process highlights TDP-43 dysfunction as a driver in disease progression. Mechanistically, the reduced nuclear levels of TDP-43 are associated with cytoplasmic aggregations and a loss of nuclear splicing functions. In a dose-dependent manner, the loss of nuclear TDP-43 results in the aberrant inclusion of exonic junctions into transcripts, often leading to transcript destabilization and degeneration through non-sense mediated decay (2–5). In neurons, the loss of specific transcripts alters the expression of proteins critical for axonal projection, which is thought to contribute to the progression of motor neuron deficits in ALS. Despite its prominent effect on neurons, TDP-43 dysfunction is observed in various cell types, including fibroblasts isolated from donors with ALS/FTD (6), pancreatic islet cells (7), and astrocytes (8–10), suggesting the possibility that its dysfunction in other cell types may also contribute to disease progression.

Early in the course of neurodegenerative diseases, increased flux across the blood brain barrier (BBB) is detected by gadolinium-based contrast agents in MRI imaging, a finding supported by elevated CSF albumin levels (11,12)(13). BBB leakage alone can exacerbate neurodegenerative changes in animal models (14). While not all the consequences of BBB leakage are fully understood, fibrin deposition has been linked to reactive changes in brain microglia (15). The BBB is a complex structure comprising endothelial cells lining the lumen of vessels, an underlying basement membrane composed of extracellular matrix proteins, and associated pericytes, astrocytes, and perivascular fibroblasts. Although each of these components contributes to the barrier, it is the endothelial cells that provide the functional barrier through tight junctions containing claudins (e.g., claudin-5) and their adapter proteins (e.g., ZO-1), along with the basal adherens junctions found throughout the vasculature (e.g. VE-cadherin). TDP-43 is highly expressed in the endothelium, but whether nuclear levels of TDP-43 contribute to BBB function is unknown.

In a parallel manuscript, we identified a loss of TDP-43 nuclear protein in an endothelial sub-population enriched in AD, and most prominently in ALS and FTD (biorxiv Ref). As this coincided with increased markers of endothelial cell activation, it led us to hypothesize that the loss of nuclear TDP-43 in endothelial cells could contribute to BBB defects in the progression of neurodegeneration. Here, we test this hypothesis, using knock-in mouse models of ALS/FTD and pan-endothelial and brain-endothelial deletion of TDP-43.

## Materials and Methods

### Generation of the Tardbp G348C mutant mouse

C57BL/6J single cell zygotes were microinjected with CRISPR/Cas9 guide RNAs flanking the Ala315 to Gly348 positioned in exon 6 of the canonical Tardbp gene and a synthesized 200nt donor dsDNA containing the following mutations (A315T (GCT->ACT), M337V (ATG->GTG) and G348C (GGC->TGC). The objective of the targeting experiment was to determine if a founder mouse containing all three clinical ALS-associated mutations in Tardbp could be recovered. This was not successful. However, founders were recovered that contained the G348C allele. Progeny derived from these founders were wild type at both A315T and M337 indicating that the donor DNA fragmented and only the G348C variant allele was established in the genome. A line was established from founder 186 and designated as JAX Strain #028435 and the G348C allele designated as Tardbp^em9Lutzy^.

### Measurement of vascular leak

Mice received a retro-orbital injection of Tomato Lectin 488 (50ul, DL-1174-1, Vector BioLabs), 3kDa Texas Red-Dextran (50uL, D3329, Thermo Fisher Scientific 10 mg/mL in PBS), EZ-Link™ Sulfo-NHS-Biotin (100uL A39256, Thermo Fisher Scientific, 1 mg/0.4 mL in PBS). After 30min circulation, mice were euthanized by CO_2_, and heparinized plasma samples were collected via cardiac puncture.

#### Direct dye measurement in the whole brain

After removing meninges, the forward half hemisphere was used for dye extraction and in 0.1% TX-100. After bead homogenization, soluble supe containing Texas Red-Dextan was measured on a fluorescent plate reader, and the signal normalized to brain weight and plasma concentration of dye.

#### Analysis of dye and NHS-biotin leak in brain section

The rear half hemisphere was fixed in 4%PFA, dehydrated in 30% sucrose and imbedded in OCT on LN_2_-cooled metal block for sectioning at 10um and 50um. Samples were directly imaged for 3kDa Texas Red-dextran, and stained with streptavidin for imaging of NHS-biotin.

### Isolation of brain endothelial cells for direct analysis of RNA

Samples with dye injection were prepared as described above. After removing meninges, one brain hemisphere was minced and digested in HBSS, and density filtered over 22% Percoll as previously described (16). The resulting pellet was resuspended in 300 µL of HBSS/BSA/glucose buffer along with the following antibodies APC-Cy7 CD45, PE CD31, PE-Cy7 ICAM-1, Alexa647 ICAM-2 at 1:100 and Live/Dead at 1:1000 (L34962, Invitrogen), for sorting of CD45neg, Lectin+, ICAM-2+ and CD31+ endothelial cells (BD FACS Aria™ III Cell Sorter, BD Biosciences).

### Isolation of murine brain endothelial cells (MBEC) for culture and in vitro treatment

Microvessels were isolated from brain tissue after removing meninges, as described (17), except that endothelial cells were allowed to grow out from microvessels on Collagen I coated plates after digestion and myelin depletion. The Growth media used was ATCC VEGF-enriched media, supplemented with Primocin, a broad-spectrum antibiotic. Cells were cultured at 3% O_2_, and 5% CO_2_, with N_2_ as balancing the gas mixture. After one day in culture, puromycin was added to the media at a concentration of 1 μg/mL. After two days, puromycin media was removed. The Cells were then sorted using ICAM2 (rat anti-mouse CD102, Clone 3C4, Catalog #105602, BioLegend) and purified (Dynabeads™ Protein G 10003D).

### Dextran Transwell Assay

MBEC were plated on a collagen-coated transwell membrane (Costar 3401) at confluency in ATCC +VEGF media. Media in the upper well was replaced 0.1mg/mL 10kDa Dextran-FITC (D1820, Invitrogen) in culture media, and sampled at the indicated timepoints from the bottom transwell for analysis (CLARIOstar Plate Reader (BMG Labtech) at a 494/518 nm wavelength). Transwell filters were collected at endpoint for analysis of cell coverage by DAPI and phalloidin.

### Immunofluorescent analysis of brain tissue and cells in culture

Brain samples, with and without dye injection described above, were fixed in 4% PFA and prepared in OCT as described above, before cutting at 10um or 50um followed by OCT removal and staining. MBEC were plated on collagen-coated 8-well chamber slides (C86024, Millipore Sigma). At ∼90% confluence, they were washed and fixed with 4% PFA. For staining of both tissue sections and cultured cells, slides were blocked/permeabilized for 15min in 2% BSA and 0.1% Triton X-100, before staining in a 1:10 dilution of block: PBS and the indicated antibodies overnight at 4C. After PBS washes, secondary antibodies (1:1000) and DAPI (D9542-1, Sigma, 1:10,000) were incubated for 2hrs at room temperature, before washing in PBS and mounting (Fluoromount, F4680, Sigma-Aldrich). To visualize the actin cytoskeleton, the cells were incubated in 5 μL of Phalloidin-FITC (5782, Bio-Techne) methanolic stock solution diluted in 200 μL of blocking solution for 1 hour in the dark at room temperature. Images for control and mutant conditions were collected in the same imaging session and with the same settings, using either a Zeiss Axioskop or Zeiss LSM-800 confocal. Raw. CZI data was obtained for analysis in ImageJ/FIJI (National Institutes of Health, Bethesda, MD, USA). Multiple fields from tissues or cells from multiple donors (>3) of each genotype were collected and data was processed in parallel.

### Antibodies

Immunofluorescence microscopy:

| Antibody | Catalog number | Provider | Dilution | Antigen retrieval |
| --- | --- | --- | --- | --- |
| anti-TDP 43 | #ab109535 | Abcam | 1:50 | Yes |
| anti-ERG | # ab196149 | Abcam | 1:100 | Yes |
| anti-Fibronectin |  | Richard O. Hynes lab | 1:200 | No |
| anti-Collagen IV | #PA1-28534 | Thermo Fisher Scientific | 1:100 | No |
| anti- Fibrinogen | #A0080 | Agilent Dako | 1:50 | Yes |
| anti-GFAP | #13-0300 | Thermo Fisher Scientific | 1:200 | Yes |
| anti-Iba1 | #ab178846 | Abcam | 1:100 | No |
| anti- VE- cadherin | #550548 | BD Biosciences | 1:100 | No |
| anti-ZO1 | #ab221547 | Abcam | 1:50 | No |
| anti-Claudin 5 | #ab131259 | Abcam | 1:50 | No |

Polyclonal antibodies against Fibronectin (297.1) were generously provided as a gift from Richard O. Hynes’s lab (Koch Institute for Integrative Cancer Research at MIT).

Flow cytometry:

| Antibody | Catalog number | Provider | Dilution |
| --- | --- | --- | --- |
| rat PE anti-mouse CD31 | #102508 | BioLegend | 1:100 |
| rat APC/Cyanine7 anti-mouse CD45 | #103116 | BioLegend | 1:100 |
| rat PE/Cyanine7 anti-mouse CD54 | #116122 | BioLegend | 1:100 |
| rat Alexa Fluor® 647 anti-mouse CD102 | #105612 | BioLegend | 1:100 |

### RNA isolation and analysis

RNA was isolated from sorted or cultured cells by RNA-easy mini or micro columns (Qiagen), with on column DNAse treatment. RNA was checked for integrity and yield before library preparation (SMART-Seq mRNA LP). Targeted sequencing was 100M 150+150 PE reads on a NovaSeq, and generally yielded more than 100M PE reads per sample. Human and mouse FASTQ were aligned (star/2.7.1a) to hg38 or mm10. For gene expression analysis the following flags were used: --outFilterMultimapNmax 20 --alignSJoverhangMin 8 --alignSJDBoverhangMin 1 --outFilterMismatchNmax 999 --alignIntronMin 10 --alignIntronMax 1000000 --alignMatesGapMax 1000000 --peOverlapNbasesMin 5. Results were assessed by Qualimap. Transcript counts (rsem/1.3.0) was performed using the following parameters --paired-end --calc-ci. For splicing analysis, FASTQ were remapped according to Leafcutter suggested parameters: --twopassMode Basic --alignIntronMin 10 --outSAMstrandField intronMotif. Output was assessed by Qualimap, before analyzing splicing according to Leafcutter protocol.

### Statistical analysis

All data collected from immunofluorescence was analyzed using the unpaired two-tailed Mann-Whitney test unless otherwise specified. All statistical analyses were conducted using GraphPad Prism software, (GraphPad Software, La Jolla, California, USA).

### Behavioral testing

#### Open Field Maze Test

Mice were tested on the Photobeam Activity System (San Diego Instruments). This system consists of a 16×16-inch clear acrylic box, and the mice were not trained prior to the trial. The mice were brought into the testing room and allowed to acclimate for 20 minutes before each trial. Each trial lasted 10 minutes, and the apparatus was cleaned before and between trials using 70% ethanol.

#### Rotarod Test

Mice were tested on a Rota-rod apparatus (ENV-574M from Med Associates Inc). Prior to the trials, mice were given one training session lasting 1 minute at 6 rpm. This was followed by three trials in which the rod’s speed was ramped up from 6 rpm to 60 rpm. Mice were brought into the testing room and allowed to acclimate for 20 minutes before the training session. Once all the mice were on the rotating rod and facing in the correct direction, the training session or trial (and subsequent ramping up of speed) was initiated. Mice were considered ‘out’ when they either fell and broke the beam at the bottom of the apparatus or when the mouse made a full revolution of the rod while hanging on, at which point the beam was tripped manually. The apparatus was cleaned before and between trials using 70% ethanol.

#### Tube Dominance Test

Mice were not trained before the trials. They were brought into the testing room and allowed to acclimate for 20 minutes before the trials. The tube used was the Tube Dominance Test (Panlab LE899M), a clear polymethyl methacrylate tube measuring 30 cm in length and 3.4 cm in diameter. This tube featured two clear gates, each located 13 cm from one end. The tube was sprayed with ethanol and dried with a paper towel before and between trials. While the mice were not specifically trained for the tube, the same mice were used for multiple trials on the same day. Mice were tested against mice of the same sex but with different genotypes from the same cage. They were placed at opposite ends of the tube and allowed to enter. Subsequent trials alternated the starting positions of the mice. Once both mice were entirely inside the tube with their noses at the gates, the gates were removed, and a timer was started. A mouse was marked as the loser when both of its feet were out of the tube, and a time was recorded. Trials that lasted more than 2 minutes were repeated at the session’s end. Trials in which a mouse turned around or passed its opponent were abandoned.

## Results

### *Tardbp*^G348C/+^ model of ALS/FTD exhibits brain barrier leak *in vivo* and impaired endothelial function

To examine brain barrier function, we injected tomato-lectin, 3kDa Texas Red-Dextran and 0.3kDa NHS-sulfo-biotin into the circulation of 10-11month old*Tardbp^G348C/+^* mice and their littermate controls. After 30 minutes of circulation, mice were euthanized and the brain extracted. The right posterior quadrant containing the cerebellum, midbrain and cortex was used for dye extraction, and analysis (Fig.1A). We found that mice *Tardbp^G348C/+^* mice exhibited increased dye leak (Fig.1B). To examine leak within specific brain vascular beds, we examined tissue sections from these mice, and found increased and diffuse staining from either the larger 3kDa Texas Red-Dextran, or the smaller 0.3kDa NHS-sulfo-biotin, detected by streptavidin 647 (Fig. 1C-F&SI Fig.1A-D). We did not observe focal leak and midbrain and cortex appeared to be similarly affected, suggesting that increased staining resulted from systemic and not localized vascular defects. Increased vascular leak did not appear to result from increased vascular density, as lectin perfused vessel density was reduced in *Tardbp^G348C/+^* mice (Fig.1G, H, SI Fig.2A&B). As BBB dysfunction is linked to astrogliosis and microgliosis (18–20), we examined astrocytes using GFAP and microglia using Iba1, and found significantly increased numbers in the*Tardbp^G348C/+^* mice, suggesting a link between brain barrier dysfunction and astrogliosis and microgliosis (Fig.1I-L&SI Fig.3A-D).

**Figure 1.**
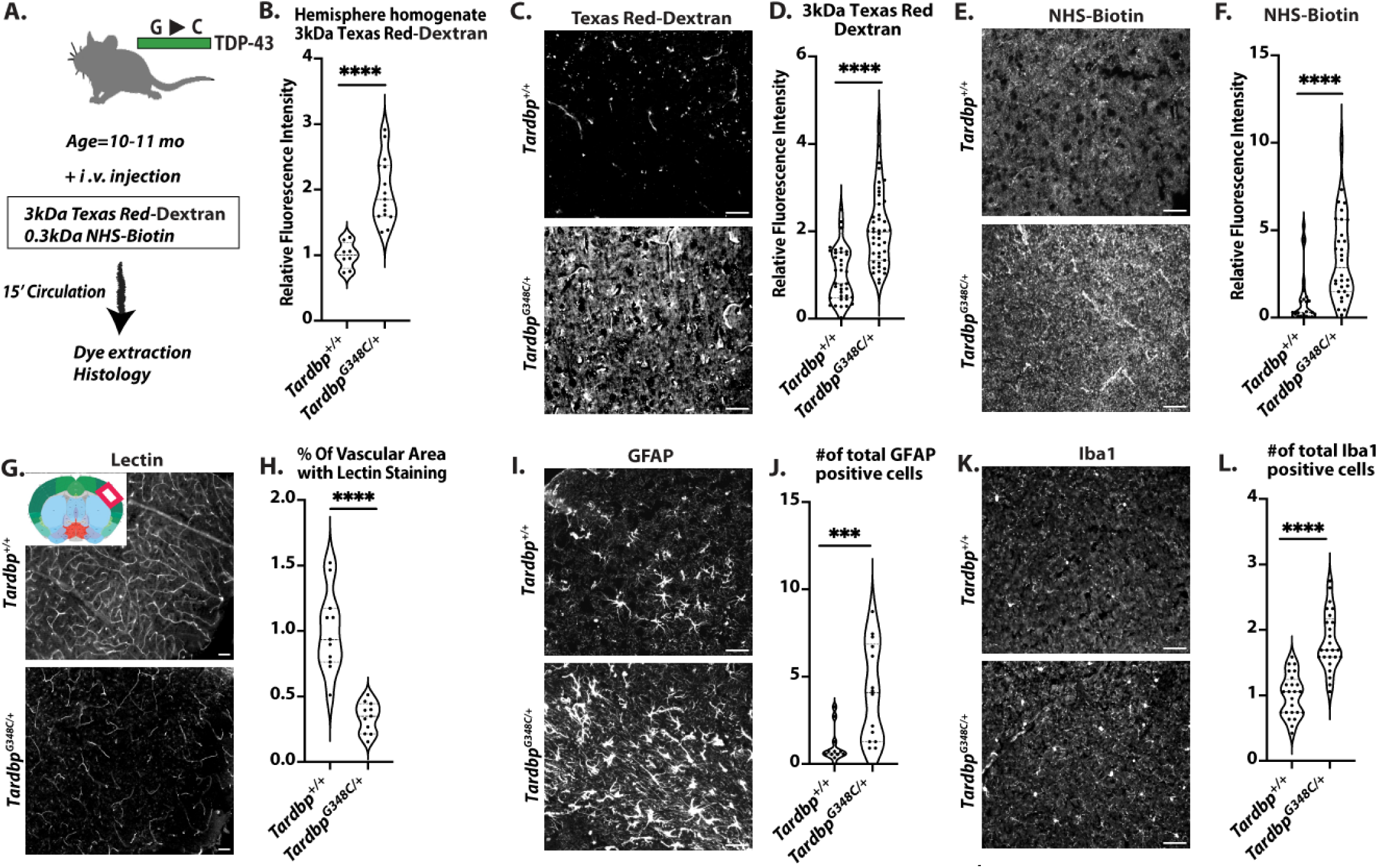
Blood brain barrier disruption and vascular sequelae in*Tardbp*^G348C/+^ mice. (A) A schematic illustration of the assay for measuring BBB permeability is presented in Figure A. (B) The quantification process involved homogenizing brain tissue samples from 10-11month-old wild-type *Tardbp^+/+^* mice (n=8) and their heterozygous littermates,*Tardbp^G348C/+^* mice (n=15), followed by measuring fluorescence intensity at 590 nm. Representative images of (C) 3kDa Texas Red-dextran leakage, (E) NHS-Biotin, (G) Tomato-lectin perfusion, (I) GFAP staining of astrocytes and (K) Iba1 staining of microglia in the mouse brain cortex reveal consistent results across*Tardbp^+/+^*mice (n=3) and *Tardbp^G348C/+^* mice (n=3). (D, F, H) Quantification of data, with each data point representing the fluorescence image intensity in one image. (J, L) Quantification of data with each data point representing the number of activated cells in an image. multiple images per mouse. Scale bars, 50 μm. Data are presented as means ± SEM. Statistical analysis was conducted using an unpaired Mann Whitney test, with significance levels indicated as follows: ***P 0.0001, ****P<0.0001.

**Figure 2.**
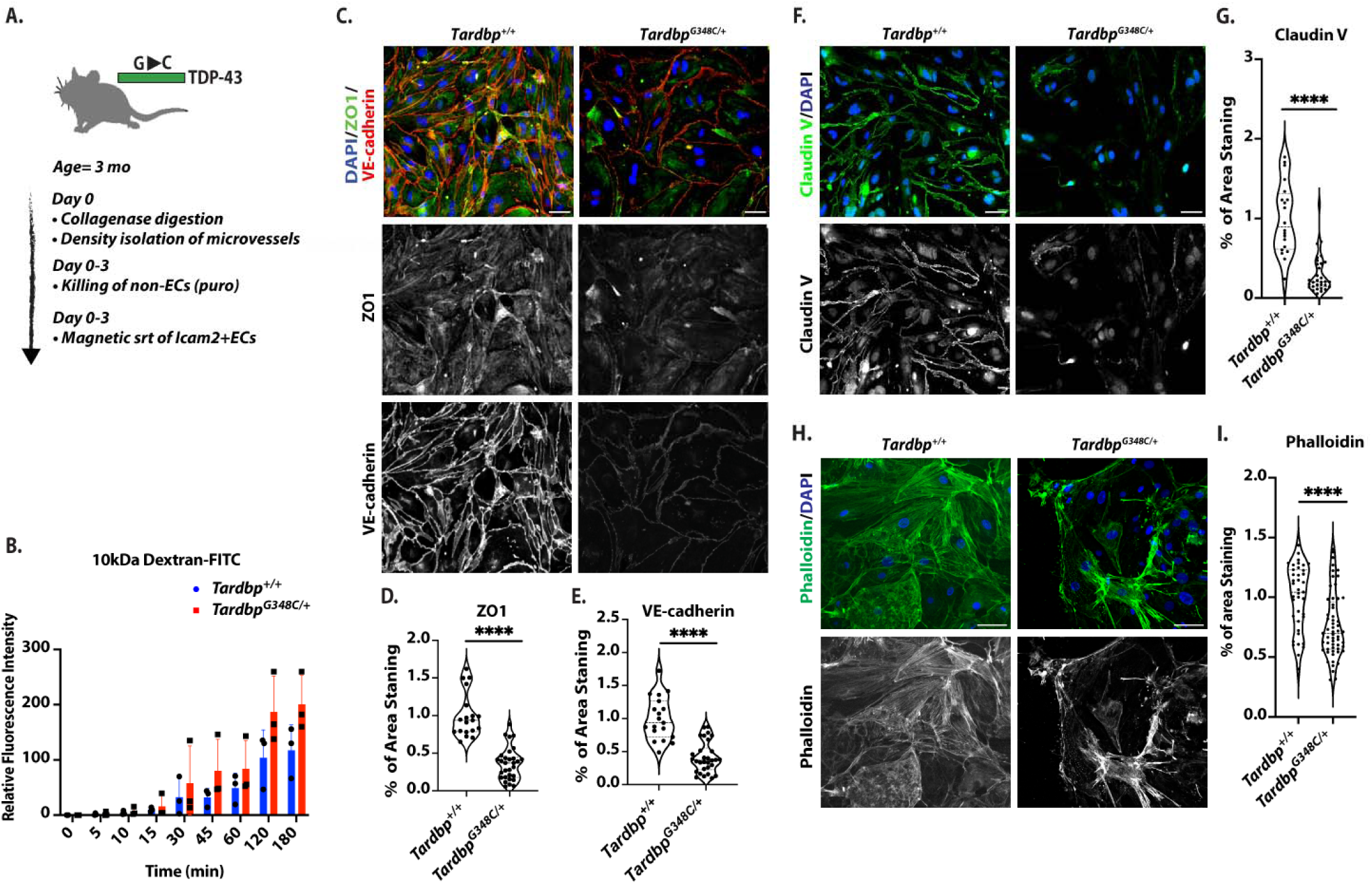
Endothelial cells isolated from *Tardbp*^G348C/+^ mice exhibited decreased levels of cell junction proteins, altered actin organization, and increased permeability. (A) Schematic illustration of the isolation and purification of ECs. (B) Passage of 10kDa FITC-Dextran dye across confluent monolayers of endothelial cells isolated from *Tardbp^+/+^*(n=3) and *Tardbp^G348C/+^* (n=3). Statistical analysis was conducted using linear regression, ****P ≤ 0.0001. (C, F, H) Representative images of mouse brain endothelial cells isolated from 3-month-old wild-type *Tardbp^+/+^*(n=6) and their heterozygous littermates,*Tardbp^G348C/+^* mice (n=6), immunostained with antibodies to the indicated proteins. (D, G, I) Quantification of data, with each data point representing the fluorescence image intensity in one image, multiple images taken of cells from each mouse. Scale bars: 50 μm. Data are presented as means ± SEM. Data are presented as means ± SEM. Statistical analysis was conducted using an unpaired Mann Whitney test, with significance levels indicated as follows: ****P ≤ 0.0001.

**Figure 3.**
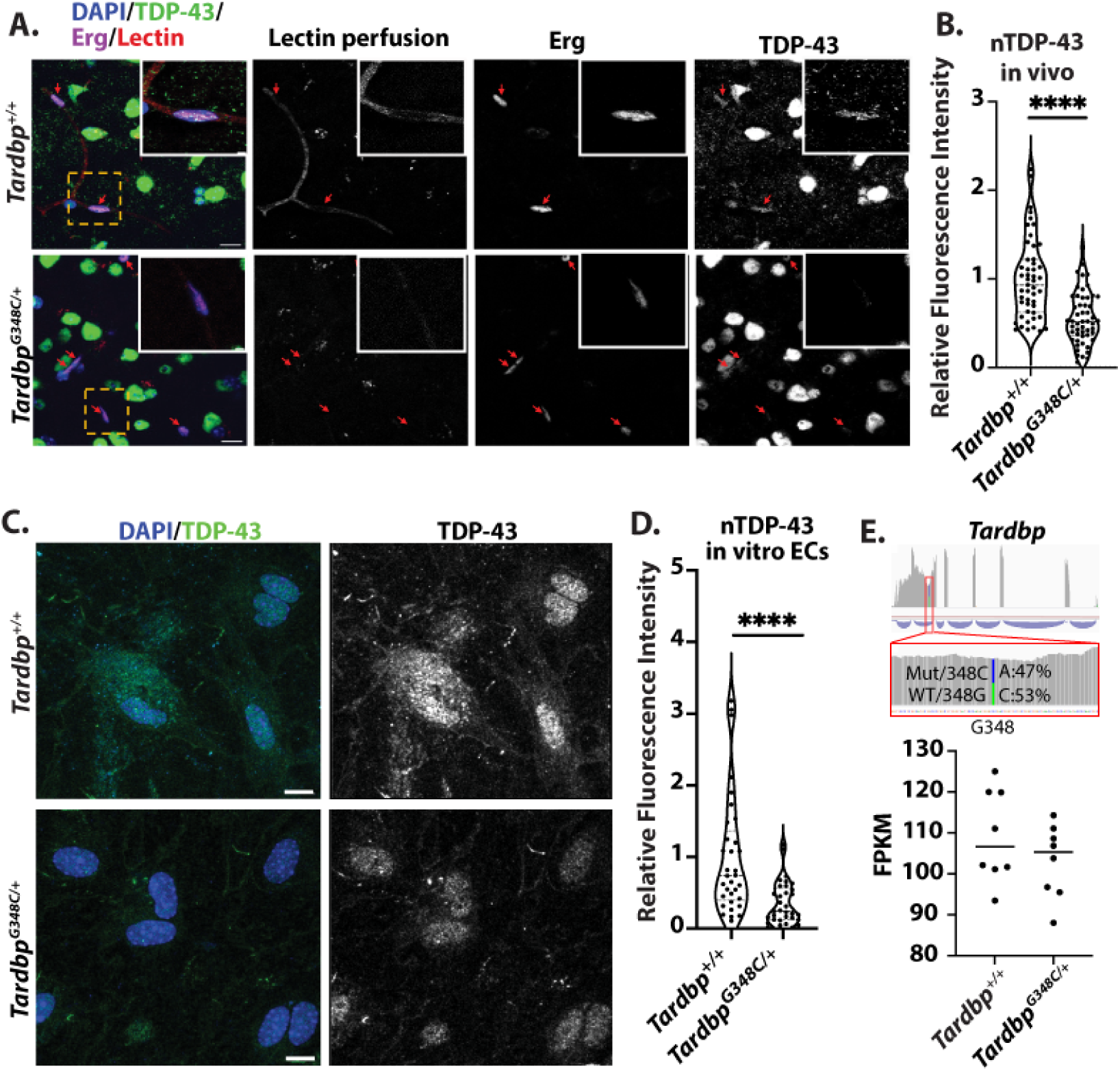
Reduced nuclear levels of TDP-43 in endothelial cells of*Tardbp*^G348C/+^ Mice. (A) Representative confocal images of mouse brain frontal cortex sections from 10-month-old wild-type*Tardbp^+/+^* (n=6) and their heterozygous littermates,*Tardbp^G348C/+^* mice (n=6), Shown are representative low magnification and high magnification (insert) immunofluorescence images for the indicated markers. Each red arrow indicates vascular endothelial nuclear TDP-43 immunostaining. (B) Quantification of TDP-43, each data point represents one vascular endothelial cell nucleus. (C) Representative confocal images of mouse brain endothelial cells isolated from 3-month-old wild-type*Tardbp^+/+^* (n=6) and their heterozygous littermates, *Tardbp^G348C/+^* mice (n=6), immunostained with antibodies against endogenous TDP-43 (green) and DAPI (blue). (D) Quantification of nuclear TDP-43 levels in isolated endothelial cells, confirming a reduction in TDP-43 in *Tardbp^G348C/+^* mice, each data point represents one endothelial cell nucleus. (E) RNA-sequencing analysis of endothelial cells derived from *Tardbp^G348C/+^* mice and littermate controls showed no significant differences in mRNA expression. Phased analysis of transcripts derived from the mutated G348C allele and the wildtype allele in the same cells revealed no differences in mRNA transcript levels. Data are presented as means ± SEM. Statistical analysis was conducted using an unpaired Mann Whitney test, with significance levels indicated as follows :****P ≤ 0.0001. (A&C) Scale bar: 50 μm.

Endothelial cells of the brain are highly specialized to provide barrier function through the expression of tight junction proteins (claudins, occludins, and JAMs) and intracellular adapters (21). Therefore, we asked whether barrier dysfunction in *Tardbp^G348C/+^* could be due to effects on the endothelium. We purified brain endothelial cells from *Tardbp^G348C/+^* mice and littermate controls, and seeded transwell filters with the same number of cells. We found that*Tardbp^G348C/+^* endothelial cells exhibited increased permeability of 10kDa FITC-dextran dye, relative to cells derived from littermate controls (Fig.2A&B). We then examined junctional proteins associated with barrier function, and observed a reduction in VE-cadherin, claudin-5 and ZO-1 (Fig.2C-G). The cytoskeleton is critical for the establishment of cell-cell junctions (22), and staining of actin by phalloidin indicated significant alterations in cellular actin (Fig.2H&I).

Thus, an ALS/FTD associated amino-acid substitution in heterozygous*Tardbp^G348C/+^* mice is sufficient to cause increased and age-associated leak across the blood brain barrier. As isolated endothelial cells exhibit defective barrier function and cell-cell junctions, the data suggests that at least some of this effect might be attributed to endothelial cell dysfunction.

### Nuclear levels of TDP-43 are reduced in the endothelial cells of*Tardbp*^G348C/+^ mice

ALS/FTD associated mutations in TDP-43 are associated with cytoplasmic accumulation and reduced nuclear levels of protein (1). To determine whether the loss of nuclear TDP-43 or cytoplasmic aggregations also occur in the brain endothelium of *Tardbp^G348C/+^* mice, similar to TDP-43 neuronal inclusions, we stained tissue sections for the protein, and examined colocalization with DAPI+ nuclei in lectin-perfused vessels in the cerebral cortex. We found that TDP-43 levels were reduced, relative to littermate controls (Fig.3A&B). To further confirm this, we examined isolated endothelial cells from cortex, and stained in culture for TDP-43. Again, we confirmed a reduction in nuclear TDP-43 (Fig.3C&D).

We then asked whether reduced nuclear TDP-43 could be a result of reduced TDP-43 mRNA in the cells. RNA-sequencing analysis *in vivo* and *in vitro* revealed no significant differences in mRNA expression in endothelial cells derived from *Tardbp^G348C/+^* mice versus their littermate controls (Fig.3E). Furthermore, phased analysis of the transcripts derived from the mutated G348C allele versus the wildtype allele in the same cells showed no differences in mRNA transcript.

Together, this data indicates the introduction of the G348C mutation in*Tardbp^G348C/+^* mice leads to a loss in nuclear levels of the protein. This is not due to a significant change in total cellular mRNA.

### Endothelial deletion of *Tardbp* elicits endothelial activation and leak in brain vasculature

A reduction in nuclear TDP-43 in *Tardbp^G348C/+^* mice, and specific defects in endothelial cells isolated from these mice suggested that endothelial cells may be sensitive to a reduction in TDP-43 levels, and that a reduction in BBB properties could be caused by a reduction in nuclear TDP-43 levels in the endothelium. To specifically address this, we generated*Cdh5(PAC)CreERT2; Tardbp^ff^* (EC-KO) mice, and *Tardbp^ff^*littermate controls. Post-natal excision of the floxed gene within the endothelium caused morbidity within 3-4 weeks (Fig.4A). The mice exhibited a range of vascular defects consistent with systemic endothelial cell activation. These included a leak of Evan’s blue dye (Fig.4B), reduced platelet counts in circulation (Figure 4C), impaired ejection fraction in the heart (Fig.4D&E) and increased fibrosis (Fig.4F).

**Figure 4.**
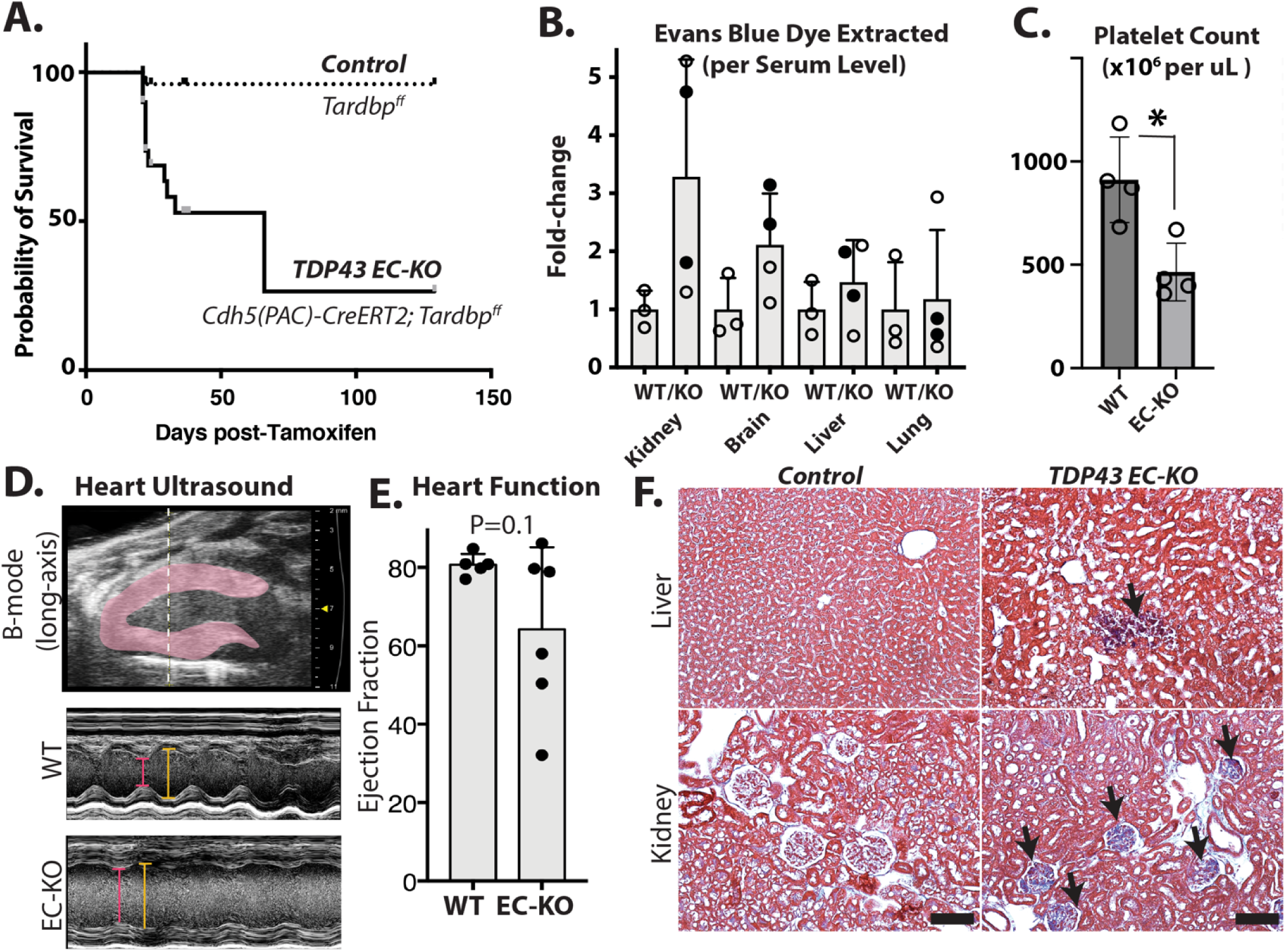
*Tardbp EC-KO* mice exhibit endothelial dysfunction and vascular leak in multiple tissues. Phenotypic analysis of TDP-43 EC-KO mice versus littermate controls at 3-4 weeks of age. (A) Mice are moribund by 3-4 weeks post-Tam treatment and gene excision. (B) Vascular leak assessed by Evans blue (EB) dye extraction (tissue EB normalized by plasma EB, presented as a fold change over the same tissue from control mice). (C) Absolute platelet count measured by flow cytometry of blood, normalized to littermate controls. (D, E) Ultrasound with quantitation, demonstrating impaired ejection fraction in EC-KO mice (n=6 EC-KO and 5 littermate controls). (F) Fibrosis was observed in trichrome-stained tissues (blue staining indicates collagen). (C) Statistical analysis was performed using an unpaired two-tailed Student’s t-test. (C&E) Statistical analysis was conducted using the Mann-Whitney test, *P<0.05.

While pan-endothelial deletion of *Tardbp* data indicated a specific requirement in the endothelium – and recapitulated endothelial permeability observed in*Tardbp^G348C/+^* mice, the rapid development of systemic vascular dysfunction precluded further analysis of brain specific functions. Therefore, we generated *Slco1c1(BAC)iCreERT2; Tardbp^ff^*(BrEC-KO) mice, and *Tardbp^ff^*littermate controls for deletion of*Tardbp* specifically within the brain endothelium. In these mice, CreER is driven by a transporter promoter with high specificity to the brain endothelium, in addition to epithelial cells of the choroid plexus (23). As expected, we observed deletion in brain endothelial cells and not in other cell-types of the brain (SI Fig.4). We found that, within a week of Tamoxifen induced gene excision in adult mice, injection of 3kDa Texas Red-Dextran accumulated at higher levels in total brain extract after inrtavascular injection, suggesting increased leak across the BBB (Fig.5A, B, SI Fig 5 A&B). Although dye leak could theoretically arise from leakage across the choroid plexus, the pattern of leak of both 3kDa Texas Red-dextran and 0.3kDa NHS-sulfo-biotin within the cortex was pervasive – and not localized to the ventricle adjacent regions, as would be expected of choroid plexus leak (Fig.5C-F). We also observed increased levels of basement membrane proteins fibronectin and collagen IV which mirrored those in the*Tardbp^G348C/+^* mice, and may be a response to BBB dysfunction (SI Fig 6&7).

**Figure 5.**
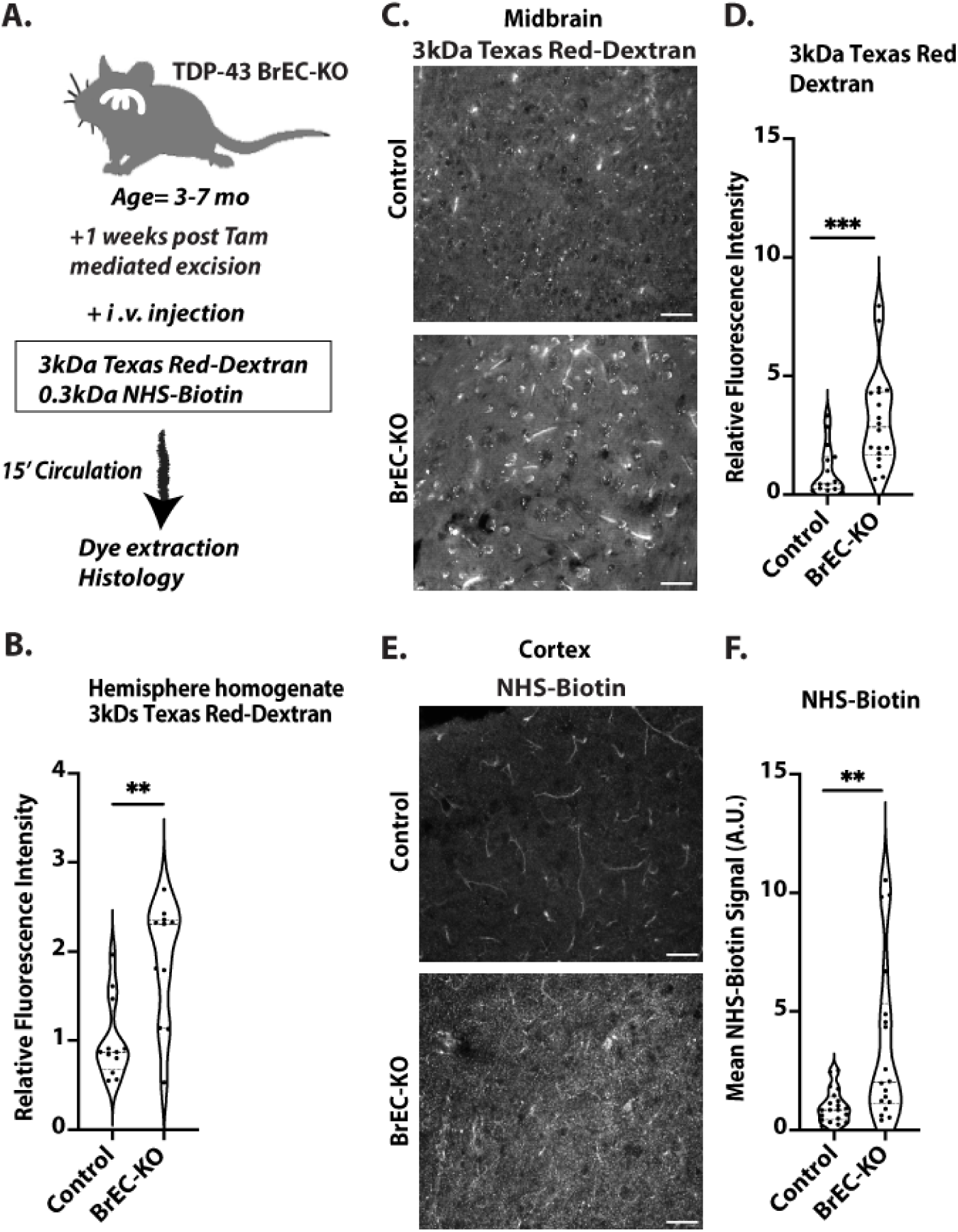
Blood brain barrier disruption in *Tardbp* BrEC-KO mice. (A) Schematic representation of the assay for measuring blood-brain barrier (BBB) permeability. (B) Measurement of 3kDa Texas Red-Dextran in homogenized brain tissue one week after Tamoxifen treatment of BrEC-KO mice (n=11) and littermate controls (n=12). **P<0.0034. (C&E) Representative images of (C) 3kDa Texas Red-Dextran leakage and (E) NHS-biotin in the cortex of 3-7-month-old mice (n=3 BrEC-KO and n=3 littermate controls). (D&F) Quantification signal, with each data point representing the fluorescence image intensity from one image, multiple images per mouse. Scale bars, 50 μm. Data are presented as means ± SEM. Statistical analysis was conducted using an unpaired Mann Whitney test, with significance levels indicated as follows: **P<0.0019, ***P<0.0002.

**Figure 6.**
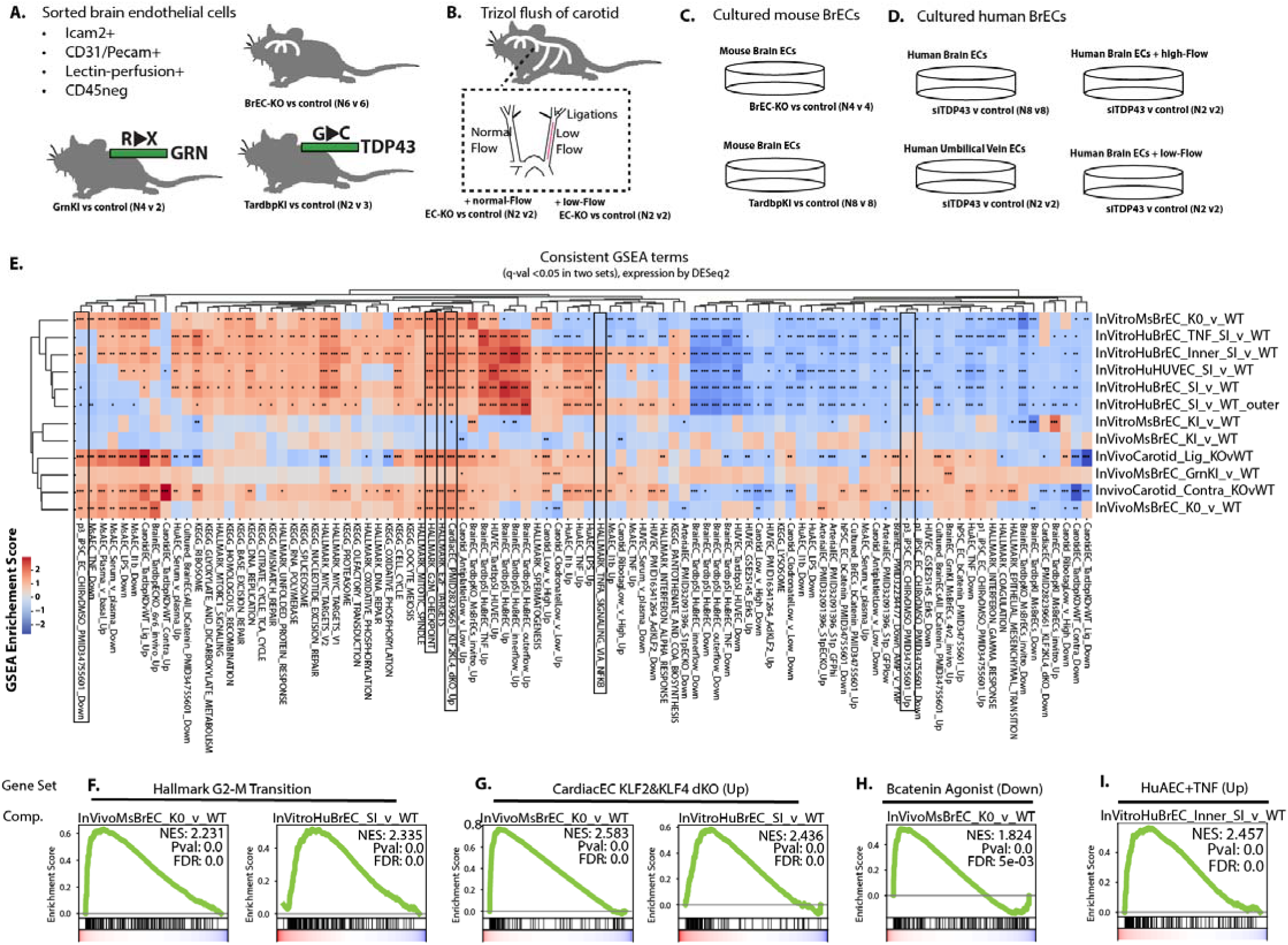
Consistent perturbations of transcriptional levels indicates core pathways targeted by partial or complete loss of nuclear TDP-43. (A-C) Outline of approach, indicating the biological replicates of each condition. (A) Shows conditions from which brain endothelial cells were sorted by flow cytometry. (B) Shows isolation procedure from carotid artery with disturbed flow. (C) Shows in vitro conditions, using primary human brain endothelial cells or purified endothelial cells isolated from BrEC-KO or KI mice. Additional details on the samples used for RNA isolation and transcriptional analysis are contained in the methods. (E) Heat map and clustering (UPGMA algorithm) of the most consistently affected KEGG and Hallmark pathways (human to mouse liftover by Biomart). Other datasets including show the top 200 transcripts up or down regulated, by p-value and after setting expression level cutoff, from a range of datasets in the literature or the lab. (F-L) Example GSEA plots for top scoring pathways. FDR value derived from GSEA analysis is shown, ***P<0.001, **P<0.01, *P<0.05.

**Figure 7.**
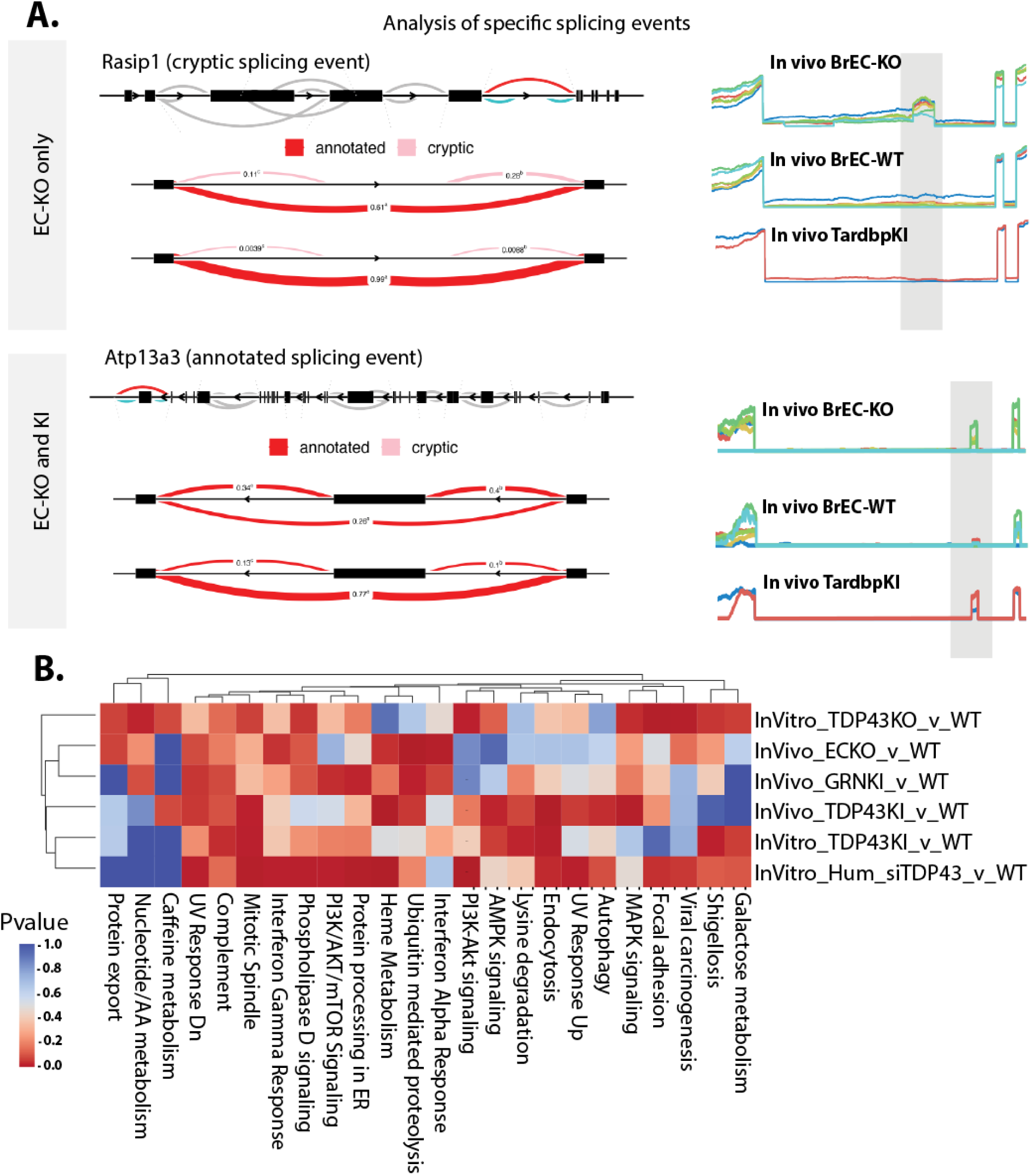
Splicing alterations associated with the loss of nuclear TDP-43 affect proteins in pathways enriched in altered mRNA transcripts. (A) Leafcutter (Leafviz) analysis of example splicing events differentially regulated in BrEC-KO cells*in vivo*, some of which are also seen in KI cells *in vivo*. Line plots to the right show read density in each of the replicates across the alternatively spliced region. (B) Heat map and clustering (UPGMA algorithm) of the most consistently affected KEGG and Hallmark pathways, and determine by GseaPy ENRICHR. (B) FDR value derived from GSEA analysis is shown.

Therefore, two independent means of endothelial *Tardbp* excision confirm a critical requirement in the endothelium, and that specific loss within brain endothelium recapitulates barrier defects observed in *Tardbp^G348C/+^* mice.

### Shared transcriptional responses between brain endothelial cells in*Tardbp*^G348C/+^*, Grn*^R493X**/+**^ and BrEC-KO mice identify pathways consistently activated by both partial and complete loss of nuclear TDP-43

Similar loss of BBB integrity in *Tardbp^G348C/+^* knockins, and in the BrEC-KO mice suggested that similar mechanisms may underlie these effects across cells with either complete or partial loss of nuclear TDP-43. To examine the transcriptional effects of TDP-43 loss, we performed RNA-sequencing analysis of acutely isolated brain endothelial cells from BrEC-KO, *Tardbp^G348C/+^, Grn*^R493X/+^, and from the carotid artery of EC-KO mice (Fig 6A&B). In addition, we performed similar analysis on cultured brain endothelial cells from BrEC-KO and *Tardbp^G348C/+^* mice under different activating conditions (no flow, laminar flow, disturbed flow, and with TNF-alpha stimulation), and the same analysis on primary human brain endothelial cells, with and without siRNA-mediated TDP-43 depletion (Fig 6C&D). By performing this analysis across a wide range of tissues with TDP-43 loss of dysfunction, we hypothesized that key transcriptional responses would be consistently altered across them.

We found pathways consistently regulated (positive enrichment or negative enrichment score, by ranked GSEA analysis), suggesting that these may be general hallmarks of TDP-43dysfunction, either partial or complete, in brain endothelial cells. These responses could be clustered into groups based on gene inclusion (Fig. 6E). As expected, knockdown and complete deletion of TDP-43 shared substantial overlap in a large set of terms, specifically related to Myc signaling, the ribosome, DNA replication, the mitotic spindle, cell cycle and G2-M checkpoint. This is consistent with a reduced proliferation of these cells in culture, and with literature which shows that TDP-43 inhibits cell-cycle at the G2-M transition (24). NFkB pathway responses were increased, indicated by analysis of either Hallmark gene sets, or the induction of these responses in human or murine arterial endothelial cells (Fig 6F-I). Wnt/b-Catenin signaling was impaired, as indicated by an increase in transcripts typically suppressed by Wnt/bCatenin signaling (Fig 6F-I).

As TDP-43 is known to regulate mRNA splicing, we then examined RNA splicing across these conditions by Leafcutter, a relatively stringent splicing analysis tool. This analysis revealed altered splicing events in several thousand mRNA transcripts including all conditions, some of which were observed primarily with complete loss of TDP-43 (EC-KO, BrEC-KO and siRNA) and some of which were observed with partial loss of TDP-43 as well (Fig 7A). Again, using the power of these datasets to identify consistently regulated pathways, we found substantial overlap with pathways affected at the transcript level. This includes mitotic spindle, focal adhesions, interferon gamma and interferon alpha, and complement (Fig 7B). However, the genes affected by mRNA splicing are different than the ones affected by total transcript levels*F*(*AK/PTK2, ITGB4, PAK1, PIK3CB* among consistently altered splicing events that could regulate focal adhesion, and *RICTOR, CCDC88A, KIF23* and *BIN1* in mitotic spindle). So, a few pathways are consistently affected by loss or depletion of TDP-43 across mouse and human endothelial cells*in vitro* and *in vivo*, via either expression or splicing.

Thus, an analysis of consistent alternative splicing patterns and gene expression in *Tardbp^G348C/+^*mice reveals transcriptional changes partially overlapping those of BrEC-KO mice.

### Chronic endothelial loss of TDP-43 results in phenotypes characteristic of Frontotemporal Dementia

*Tardbp* knock-in mouse models of ALS/FTD, like*Tardbp^G348C/+^*, may affect multiple cell types within the brain. To determine whether hallmarks of FTD might be linked to TDP-43 loss from the endothelium, we examined these hallmarks 11 months after post-natal excision of TDP-43.

First, we stained for fibrin, which is deposited near compromised brain vessels and contributes directly to neuronal damage (20,25,26). We found increased fibrin staining around the vessels in the cerebral cortex and midbrain (Fig.8A, B, SI Fig.8A&B), suggesting chronic defects in BBB function. Notably, this was associated with alterations in basement membrane components, as we observed increased levels of both fibronectin and collagen IV in BrEC-KO mice, resembling KI mice (SI Fig.6&7). Fibrin deposition has been linked to microglial activation, a common response observed in clinical FTD and animal models of the disease (27–29). Therefore, we stained sections from BrEC-KO and littermate controls with Ibα1. We found increased numbers of microglia in BrEC-KO, consistent with increased microglial activation (Fig.8C&D, P<0.0001, SI Fig.9A&B). Notably, we used a bona fide mouse model of FTD, based on a knock-in mutation in progranulin, *Grn*^R493X/+^ (27), and found that the increase in microglial numbers in the BrEC-KO closely paralleled this mouse model (Fig.8E, F SI Fig.9C&D). Astrogliosis has also been observed in FTD, where it occurs prior to neuronal loss. Therefore, we stained and counted GFAP+ cells. We found a substantial increase in the number of astrocytes in BrEC-KO mice and*Grn*^R493X/+^ mice relative to littermate controls (Fig.8G-J SI Fig.9E&H). Thus, the combination of increased perivascular fibrin deposition and microglial activation indicates chronic damage to the BBB resulting in immune activation in the brain.

**Figure 8:**
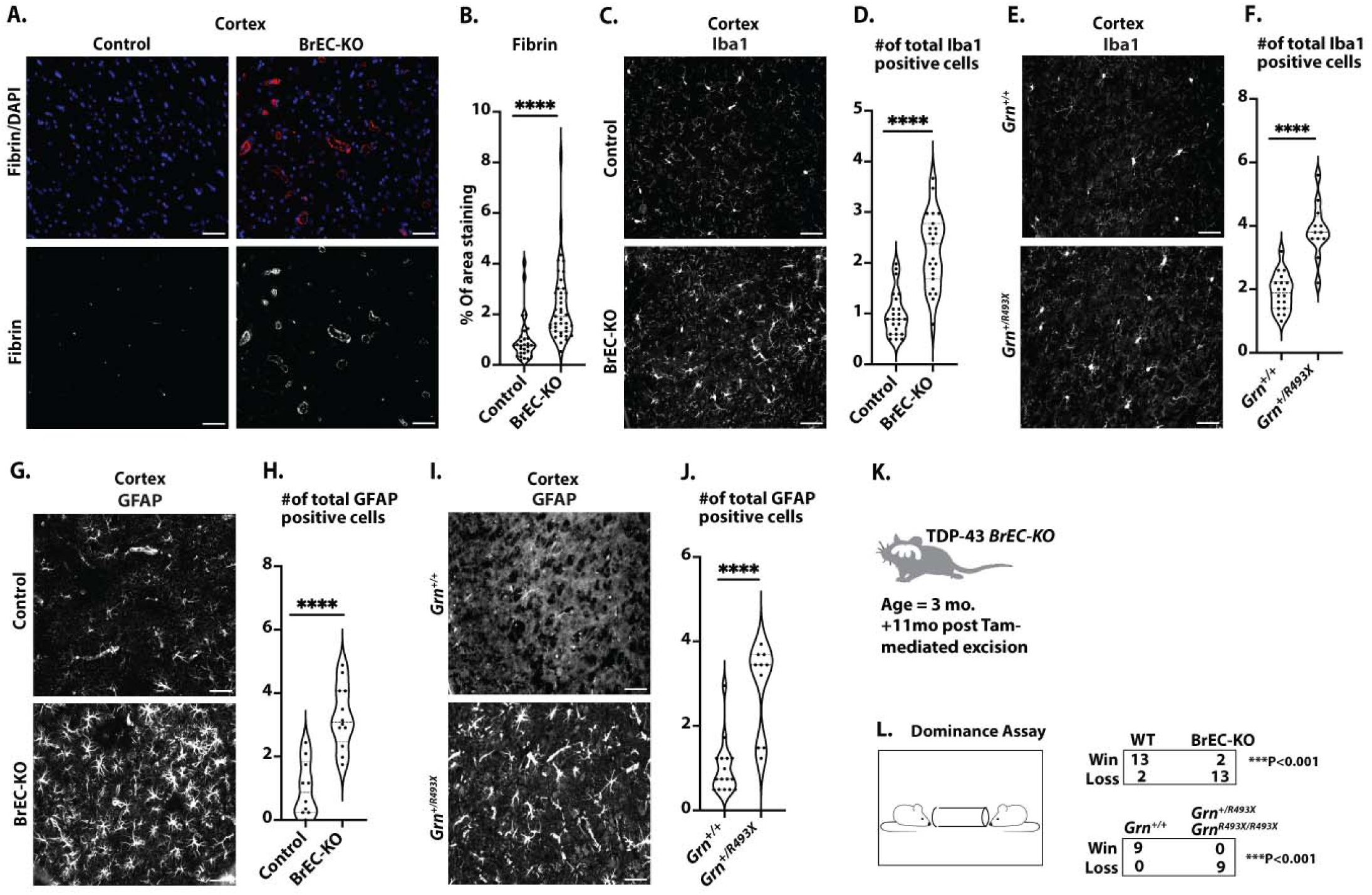
Pathological and behavioral consequences of chronic endothelial TDP-43 loss. (A) Representative immunofluorescence images of fibrin deposition in mouse brain frontal cortex sections from 8-11-month-old mice (n=3 BrEC-KO and n=3 littermate controls) are shown. (C, E) Iba1 staining of microglia in the mouse brain cortex reveals consistent results across n=3 BrEC-KO and n=3 littermate controls mice (n=3), as well as n=3 *Grn^R493X/+^* and n=3 littermate controls mice (n=3) and (G, I) GFAP staining of astrocytes reveals a substantial increase in astrocyte numbers, resembling astrogliosis observed in FTD. (K, L) Behavioral Testing: Tube dominance test results for BrEC-KO mice and*Grn^R493X/+^* mice, showing a high “loss” percentage in both models. (B) Quantification of data, with each data point representing the fluorescence image intensity in one image. (D, F, H, J) Quantification of data with each data point representing the number of activated cells in an image. multiple images per mouse. Scale bars, 50 μm. Data are presented as means ± SEM. Statistical analysis was conducted using an unpaired Mann Whitney test, with significance levels indicated as follows: ****P<0.0001.

Of the behavioral tests most reliably associated mouse models of FTD, a tube dominance test has accurately segregated FTD models from controls (30). Therefore, we examined both BrEC-KO mice and *Grn*^R493X/+^ mice, along with their littermate controls in this assay. As expected, in cage-mate matches, the*Grn*^R493X/+^ mice backed away from their wild-type controls, with a 100% “loss” percentage for*Grn*^R493X/+^ mice (Fig.8K&L). We observed a strikingly similar phenotype in the BrEC-KO mice, with an 87% “loss” percentage for these mice (Fig.8K&L). We observed no differences in either basal activity in open field test (SI Fig.11). The*Grn*^R493X/+^ mice also show no defects in these assays, as expected. Therefore, in addition to pathological markers of cortical damage in FTD (fibrin deposition and microglial activation) and reduced microvascular perfusion (SI Fig 9&10), the chronic BrEC-KO mice exhibit behavior consistent with a gold-standard model of FTD.

## Discussion

Here, we show that TDP-43 has a critical function in the maintenance of the blood brain barrier. A single mutated allele of *Tardbp*, harboring a point mutation found in patients with ALS/FTD, is sufficient to cause age dependent disruption of BBB *in vivo*. This coincides with a reduction in nuclear levels of TDP-43 and cell-cell tight and adherens junctions. A reduction in nuclear levels of TDP-43 within the brain endothelium acutely results in a loss of barrier, and chronically, over several months, leads to hallmarks of FTD in the cerebral cortex, including astrogliosis and microglial activation and behavioral defects resembling a gold-standard model of FTD, based on a loss-of-function mutation in*Grn*. Notably, several key processes are affected at the RNA level in both mouse and human cells harboring disease associated mutations linked to nuclear TDP-43 loss, or direct silencing of the gene, suggesting a likely overlap between murine and human endothelial functions for TDP-43.

### TDP-43 and blood brain barrier maintenance

Despite the early focus on TDP-43 function in neurons as a mediator of ALS/FTD, it is now appreciated that this fairly ubiquitous splice factor is likely to affect the function of most cells, including fibroblasts, astrocytes and even cells of the pancreatic islet (6–10,31). It has also become clear that the transcriptional effects of TDP-43 dysfunction exert cell and species-specific effects on RNA splicing. Most notably, cryptic exons, which appear upon loss of TDP-43, show little overlap between stem cells, neurons, and muscle (32). Here, we reveal a direct effect on the BBB in animal models that occurs, in part, through a loss of cell-cell junction proteins. Our data is consistent with prior work showing that the loss of TDP-43 orthologs in zebrafish led to specific vascular defects (33,34). Although this work did not examine endothelial or brain barrier function, it did identify defects in matrix interaction, revealed by increased endothelial expression of FN1 and Integrin α4. These defects were responsible, at least in part, for cell migration defects observed in the fish. In mouse models, In an animal model, loss of*Grn* – which we show here is associated with a loss of TDP-43 in the endothelium – is linked to increase vascular permeability (35). Overexpression of TDP-43 throughout the brain via AAV injection leads to increased extranuclear TDP-43 inclusions and also increased BBB leak, but it was unclear whether the effects were through endothelial cells*in vivo*, although it could affect some of the same junctional proteins we saw altered*in vitro* (36,37). By revealing that ALS/FTD mutations drive cell-autonomous effects in the endothelium and that specific targeting of TDP-43 in brain endothelial cells *in vivo* results in rapid increases in BBB permeability, our data implicates this pathway in BBB disruption in ALS, FTD and perhaps also the many other situations in which both TDP-43 and the BBB are disrupted, including traumatic brain injury, Alzheimer’s disease and Huntington’s disease.

### Hallmarks of FTD with loss of TDP-43 and blood brain barrier

Frontal temporal lobe dementia is defined by a loss of neuron function in the frontal temporal lobe, broadly diminished social conduct and foresight, and also reduced language and speech. Rodent models of FTD are imperfect representations of these behaviors, but one of the most consistent behavioral phenotypes in a mouse model of FTD is an age dependent impairment of social interaction in a tube dominance assay (30). Here, we show that BBB defects caused by endothelial specific loss of TDP-43 result in chronic but not acute defects in this assay. Although we did not observe clear motor neuron defects, for example in the rotarod assay, it is possible that the Slco1a1-CreER approach we used failed to target endothelium near the motor neurons well. Analysis of mT/mG labeled CreER activity in the mouse identified Cre-mediated GFP signal mainly in the cerebral cortex, the area affected in FTD.

In BrEC-KO mice, behavioral defects are likely a consequence of chronic leak of multiple plasma components, and fibrinogen in particular, across the BBB – leading to activation of microglia and astrocytes, and damage to neurons (15). Acute vascular leak has been linked to cognitive defects, for example following post-natal deletion of the S1P receptor in the endothelium (38), and after acute astrocyte or pericyte deletion (18,39). Alterations in astrocyte TDP-43, acutely induced by overexpression of a transgene in mice, leads to memory impairment – but it is not clear whether this was associated with effects on the BBB(10). *Grn* mutant mice, which we show here have reduced nuclear levels of endothelial TDP-43, exhibit both barrier defects and defects in tube dominance assay (30,35). Our data shows that endothelial loss of TDP-43 is sufficient to cause fibrin deposition, significant BBB leak, and behavioral defects resembling FTD models.

### Cytoskeletal alterations as a basis of BBB defects

Our data reveal specific pathways associated with microtubules and ECM-integrin interactions which were altered across a range of models, including*Tardbp* deletion *in vivo* and *in vitro*, siRNA in human cells *in vitro*, and FTD-associated mutations linked to reduced nuclear levels of TDP-43. Nuclear levels of TDP-43 can affect RNA splicing in a dosage-sensitive manner in neurons (2), and often in a species-dependent manner (4). Low levels of TDP-43 depletion affect sensitive targets, and complete depletion of TDP-43 uncovers the most sensitive targets. In our data, FTD mutations in*Tardbp* or *Grn* result in only a partial loss of nuclear function. Nevertheless, we see similar pathways affected by both partial and complete loss of*Tardbp*, for example, centrosome, DNA replication, proteasome and TNF/NFkB signaling. The DNA replication defects are consistent with DNA replication defects in cells, resulting in part from the development of R-loops during replication (40). Interestingly, recent data is supporting an association with TDP-43 and the centrosome, although its function there is not yet clear (41). There is also literature to support an interaction between NFkB pathway transcription factor RELA/p65 and TDP-43, however – at least in microglia – the interaction appeared to be positive in leading to downstream NFkB target gene stimulation, opposite to what we see in the endothelium (42). That partial and complete loss of TDP-43 leads to similar pathway effects suggests that these pathways in the endothelium are highly sensitive to alterations in TDP-43 function. Even a partial loss of function with age, or as a result of mutations affecting TDP-43 or its regulator proteins (C9orf72, *GRN*) would likely lead to endothelial cell dysfunction.

In summary, our data demonstrate that endothelial cells are highly sensitive to the levels of TDP-43. As reduced nuclear levels of TDP-43 are caused by ALS/FTD-associated mutations and endothelial barrier defects, we propose the TDP-43 alterations within the endothelium contribute to BBB permeability and cognitive dysfunction in ALS/FTD, and perhaps other diseases where TDP-43 dysfunction has been reported.

## Acknowledgements

This work was supported by UConn Health startup funds from the School of Medicine and Department of Cell Biology, Center for Vascular Biology and Calhoun Cardiology Center, American Heart Association Innovative Project Award 19IPLOI34770151 (to P.A.M.); NIH National Heart, Lung, and Blood Institute Grants K99/R00-HL125727 and RF1-NS117449 (to P.A.M); American Heart Association Predoctoral award (to O.M.F.O.); and U54 grant U54 OD020351 (to C.L.).We recognize contributions from The Jackson Laboratory Genome Technologies Services for technical assistance and consultation. The Jackson Laboratory scientific services are supported in part through the National Cancer Institute’s Cancer Core Grant P30CA034196. We are grateful for the consistent help from Vijendar Singh and the staff at the Computational Biology Core at UConn Health in providing input on computational approaches and installing packages on the cluster. Finally, we appreciate the help of the Bernard Cook, Science Writer and Illustrator in the Dean’s Office at the School of Medicine at UConn Health, who prepared illustrations for this work, and for other members of the lab, especially Omar MF Omar, who provided consistent input into the research described here.

## Author contributions

Ashok Cheemala (AC), Amy L. Kimble (ALK), Jordan D. Tyburski (JDT), Nathan K. Leclair (NKL), Aamir R. Zuberi (ARZ), Melissa Murphy (MM), Evan R. Jellison (ERJ), Bo Reese (BR), Xiangyou Hu (XH), Catherine M. Lutz (CML), Riqiang Yan (RY), Patrick A. Murphy (PAM) AC and PAM designed the research; AC, ALK, MM, NKL and PAM performed the research; The mouse colony was managed, and genotyping was conducted by AC, ALK, and JDT; CML and ARZ generated the Tardbp G348C mouse; ALK and PAM generated the Tardbp BrEC-KO and EC-KO mice. AC and ERJ conducted FACS analysis; BR generated the RNA-Seq data; ALK and XH performed behavioral assays; AC and PAM analyzed the data and wrote the manuscript with input from RY.

Correspondence to Patrick A. Murphy.

---

**SI Figure 1.**
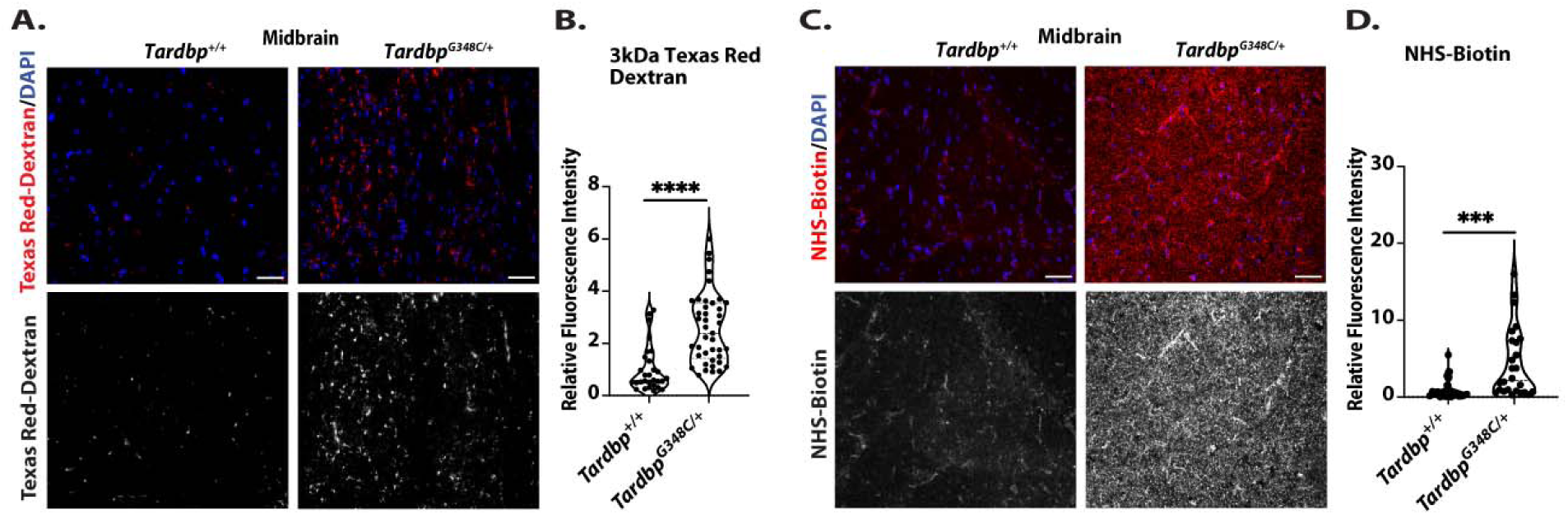
Assessment of Blood-Brain Barrier Permeability. **(A)** The representative immunofluorescence images of 3kDa Texas Red-dextran leakage in mouse midbrain sections from 10-11-month-old mice (n=3 *Tardbp^+/+^* and n=3 *Tardbp^G348C/+^*^)^ are shown. (C) NHS-Biotin. (B, D) Quantification of data, with each data point representing the fluorescence image intensity in one image. multiple images per mouse. Scale bars, 50 μm. Data are presented as means ± SEM. Statistical analysis was conducted using an unpaired Mann Whitney test, with significance levels indicated as follows: ***P 0.0001, ****P<0.0001.

**SI Figure 2.**
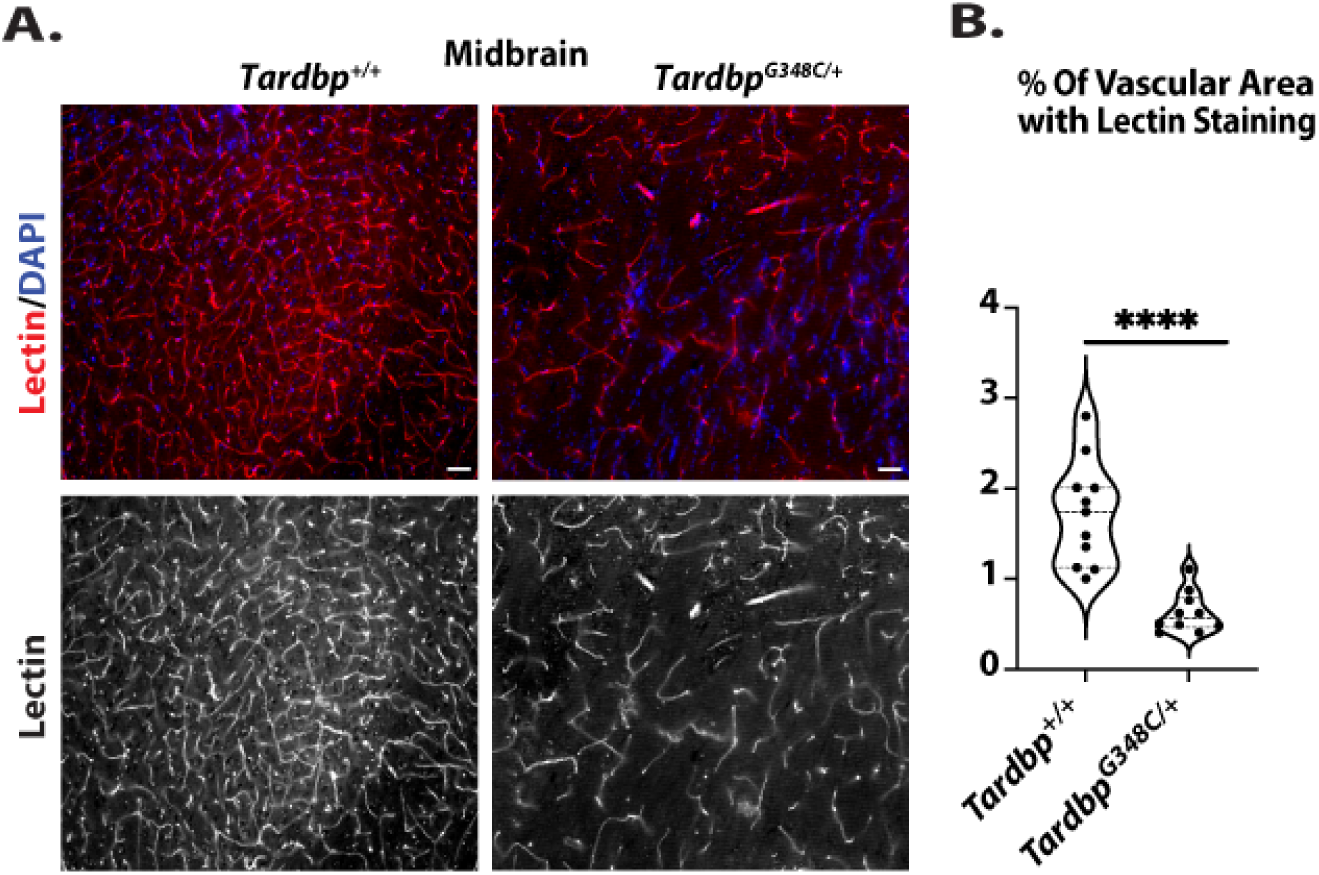
Evaluation of Tomato-Lectin Perfusion. **(A)** The representative immunofluorescence images of Tomato-lectin perfusion in mouse midbrain sections from 10-11-month-old mice (n=3*Tardbp^+/+^*and n=3 *Tardbp^G348C/+^*^)^ are shown. (B) Quantification of data, with each data point representing the fluorescence image intensity in one image. Multiple images per mouse. Scale bars, 50 μm. Data are presented as means ± SEM. Statistical analysis was conducted using an unpaired Mann Whitney test, with significance levels indicated as follows: ****P<0.0001.

**SI Figure 3.**
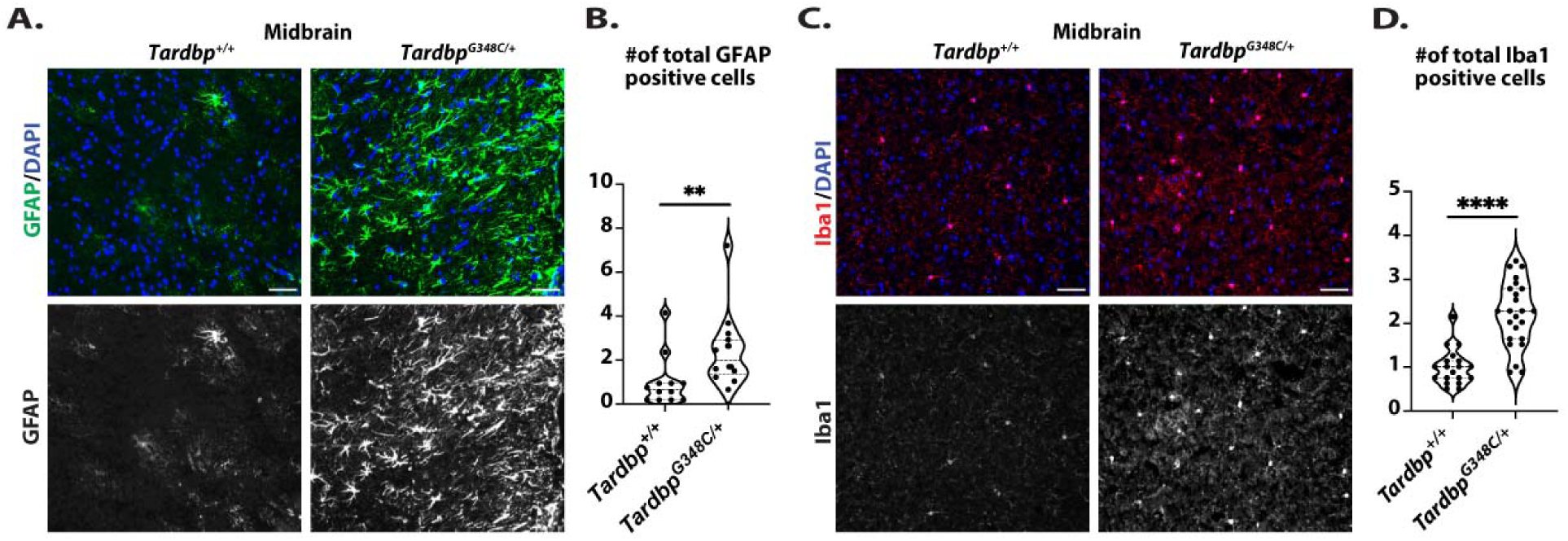
Assessment of Astrocyte (GFAP) and Microglia (Iba1) Activation. **(A)**The representative immunofluorescence images of GFAP staining of astrocytes, and (C) Iba1 staining of microglia in the mouse midbrain reveal consistent results across*Tardbp^+/+^* mice (n=3) and *Tardbp^G348C/+^* mice (n=3). (B, D) Quantification of data with each data point representing the number of actvi ated cells in an image. multiple images per mouse. Scale bars, 50 μm. Data are presented as means ± SEM. Statistical analysis was conducted using an unpaired Mann Whitney test, with significance levels indicated as follows: **P 0.0023, ****P<0.0001.

**SI Figure 4.**
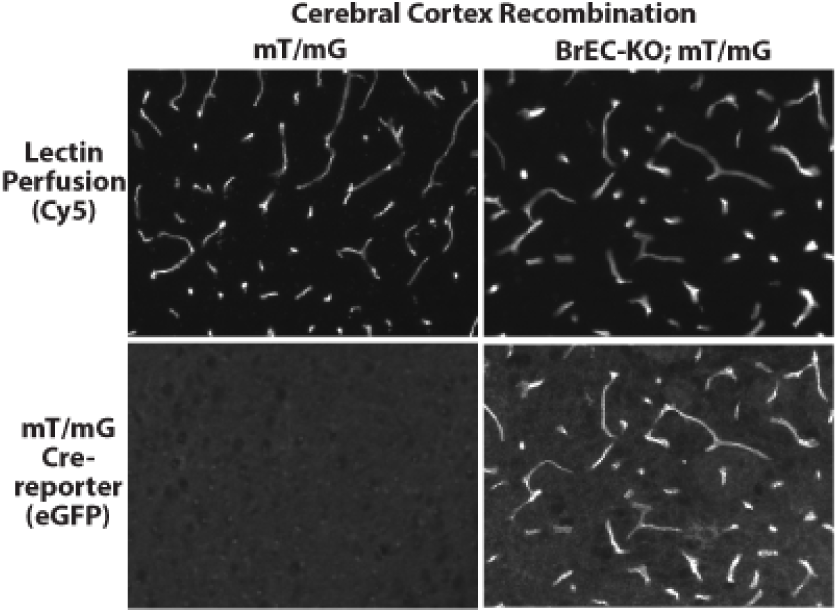
Slco1c1-CreERT2 is active in cortical vasculature. (A) BrEC-KO mice with mT/mG Cre reporter were treated with Tamoxifen and collected with Dylight649 lectin perfusion 12 weeks later. eGFP indicates Cre activity in cortical vessels.

**SI Figure 5.**
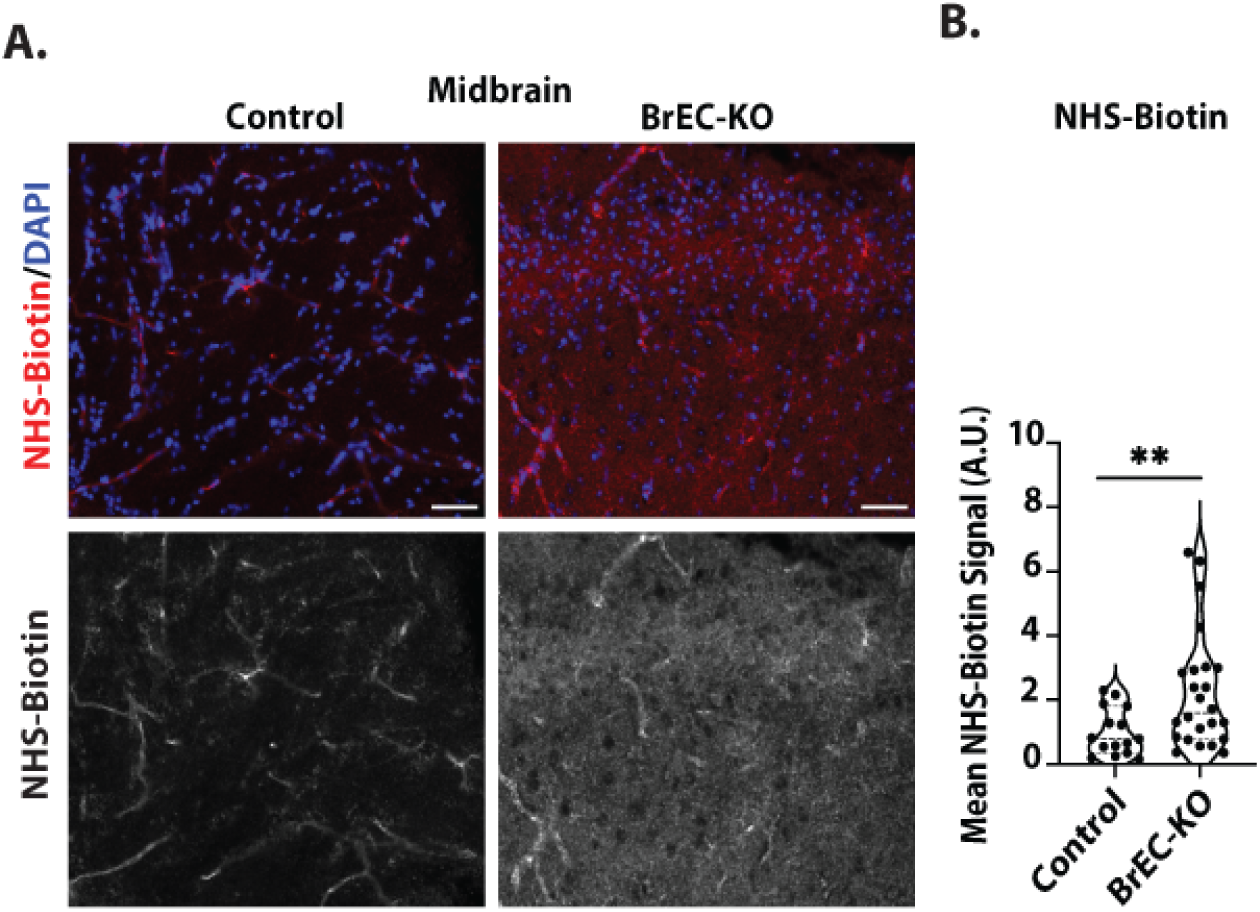
Blood brain barrier disruption in*Tardbp* BrEC-KO, midbrain. (A) The representative immunofluorescence images of NHS-biotin leakage in mouse midbrain sections from 3-7-month-old mice (n=3 BrEC-KO and n=3 littermate controls) are shown. B) Quantification signal, with each data point representing the fluorescence image intensity from one image, multiple images per mouse. Scale bars, 50 μm. Data is presented as means ± SEM. Statistical analysis was performed using an unpaired two-tailed Mann Whitney test, with significance levels indicated as follows: \ast \ast PL0.0096.

**SI Figure 6.**
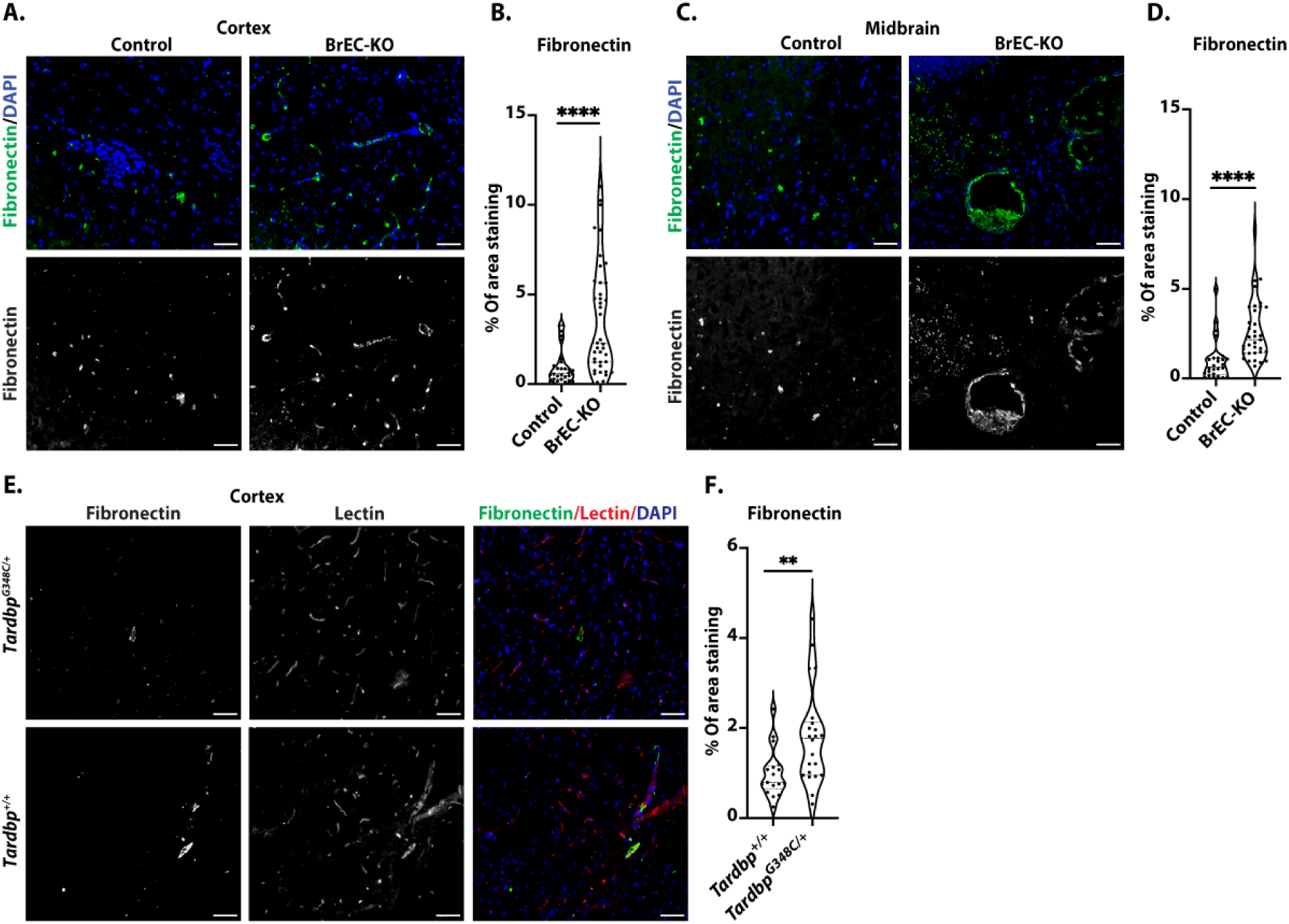
Increased Fibronectin Expression in*Tardbp* BrEC-KO and *Tardbp*^G348C/+^ mice. (A) Representative immunofluorescence images of fibronectin in mouse cortex and (C) midbrain sections from 8-11-month-old mice (n=3 BrEC-KO and n=3 littermate controls) are presented. Additionally, (E) representative immunofluorescence images of fibronectin in mouse cortex sections from 10-11-month-old mice (n=3 *Tardbp^+/+^* and n=3 *Tardbp^G348C/+^*^)^ are shown. (B, D, F) Quantification signal, with each data point representing the fluorescence image intensity from one image, multiple images per mouse. Scale bars, 50 μm. Data is presented as means ± SEM. Statistical analysis was performed using an unpaired two-tailed Mann Whitney test, with significance levels indicated as follows: **PL0.0044, ****P<0.0001.

**SI Figure 7.**
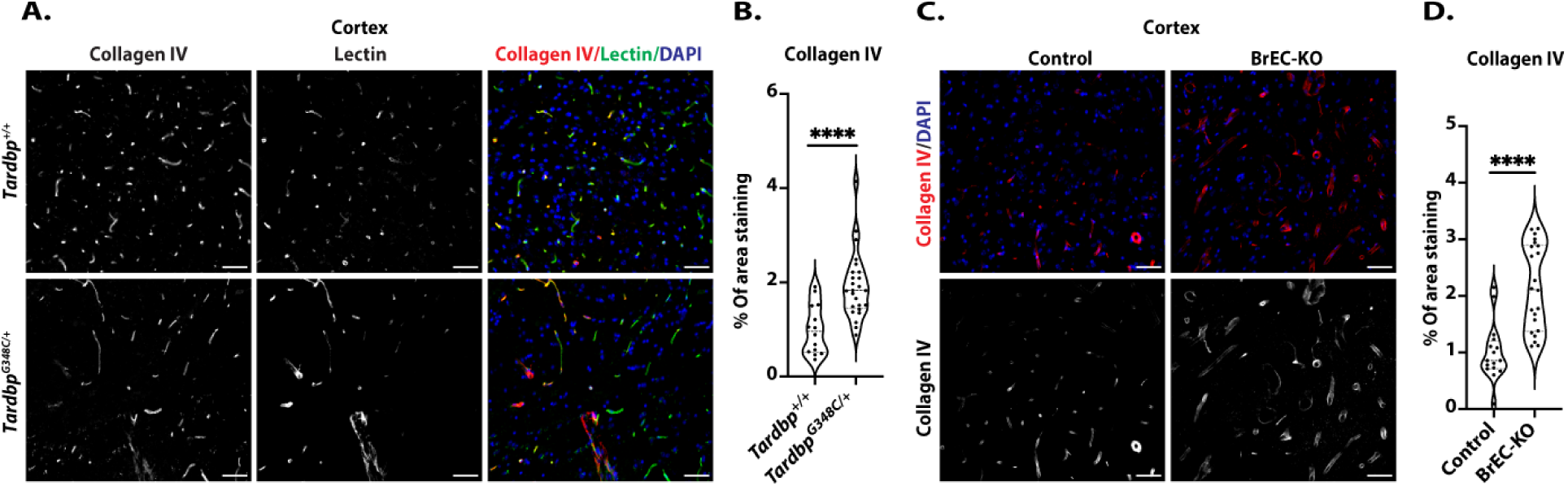
Increased Collagen IV Expression in *Tardbp* BrEC-KO and *Tardbp*^G348C/+^ mice. (A) Representative immunofluorescence images of Collagen IV in mouse cortex sections from 10-11-month-old mice (n=3 *Tardbp^+/+^* and n=3 *Tardbp^G348C/+^*) are presented. Additionally, (C) representative immunofluorescence images of Collagen IV in mouse cortex sections from 8-11-month-old mice (n=3 BrEC-KO and n=3 littermate controls) are shown. (B&D) Quantification signal, with each data point representing the fluorescence image intensity from one image, multiple images per mouse. Scale bars, 50 μm. Data is presented as means ± SEM. Statistical analysis was performed using an unpaired two-tailed Mann Whitney test, with significance levels indicated as follows: ****P<0.0001.

**SI Figure 8.**
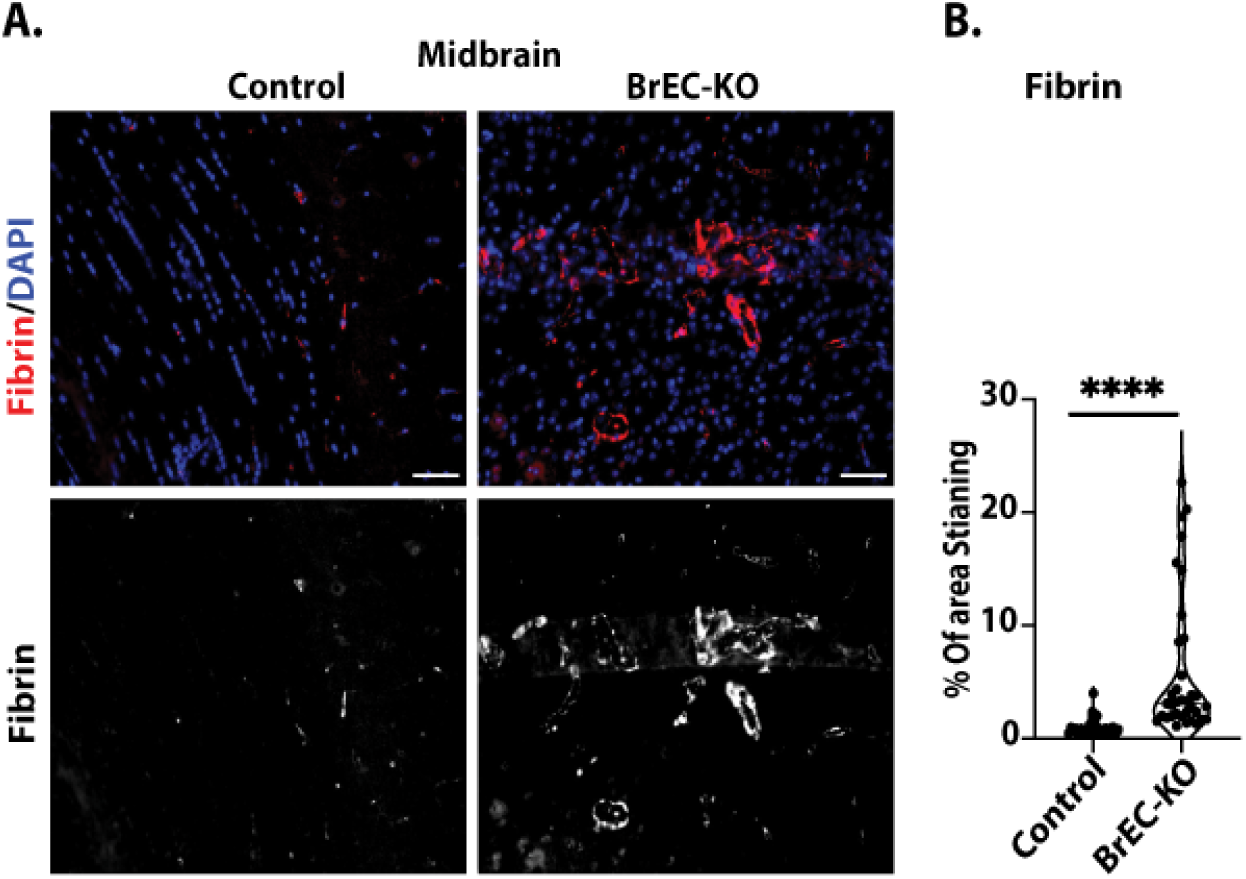
Fibrin Deposition in BrEC-KO Mouse Midbrain. (A) Representative immunofluorescence images of fibrin deposition in mouse midbrain sections from 8-11-month-old mice (n=3 BrEC-KO and n=3 littermate controls) are shown. (B) Quantification signal, with each data point representing the fluorescence image intensity from one image, multiple images per mouse. Scale bars, 50 μm.Data is presented as means ± SEM. Statistical analysis was performed using an unpaired two-tailed Mann Whitney test, with significance levels indicated as follows: ****P<0.0001.

**SI Figure 9.**
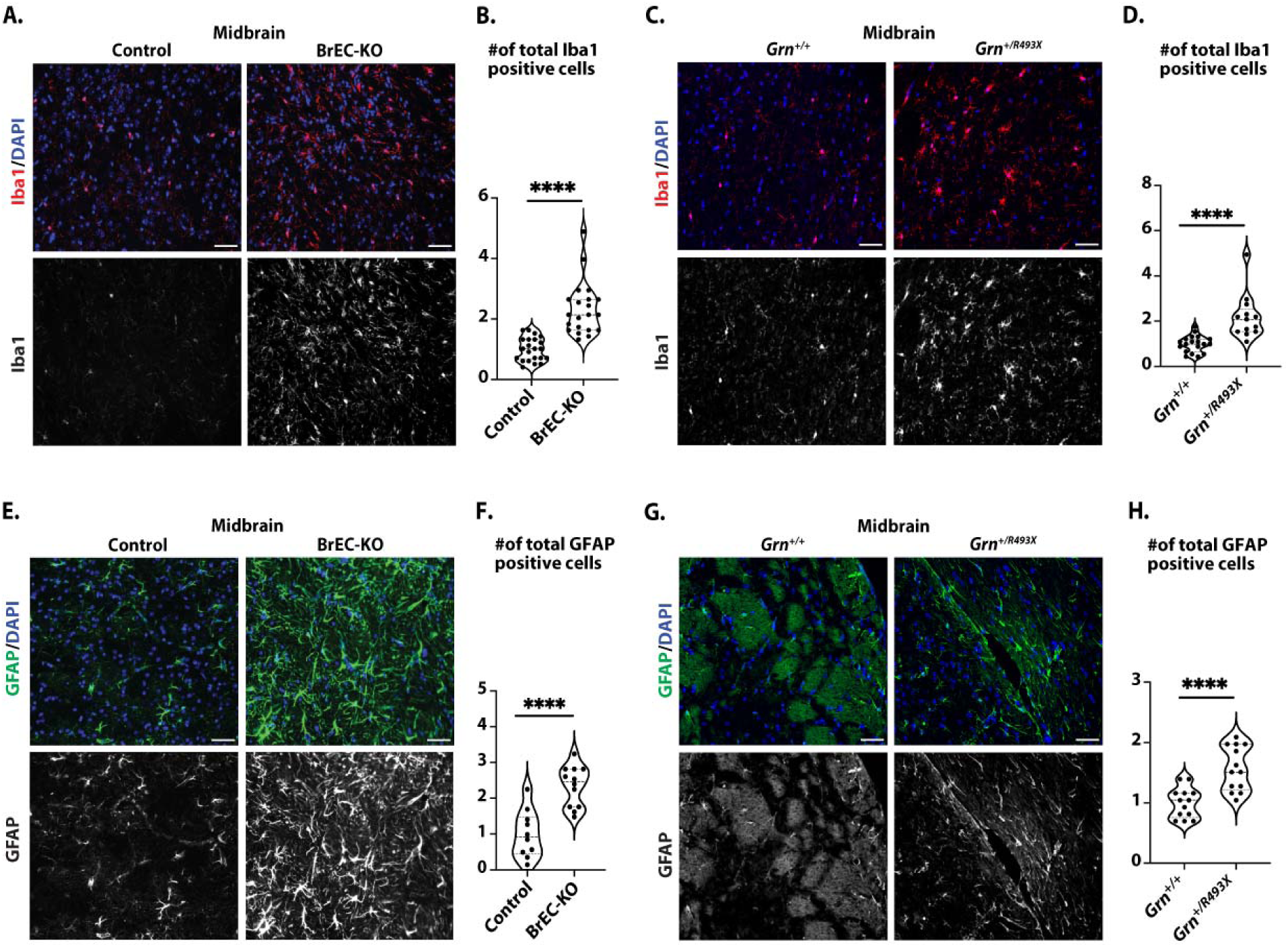
Assessment of Astrocyte (GFAP) and Microglia (Iba1) Activation. (A, C) Representative immunofluorescence images of Iba1 staining of microglia in the mouse midbrain reveal consistent results across (n=3 BrEC-KO and littermate controls mice (n=3), as well as n=3*Grn^R493X/+^* and littermate controls mice (n=3) and (E, G) GFAP staining of astrocytes reveals a substantial increase in astrocyte numbers, resembling astrogliosis observed in FTD. (B, D, F, H) Quantification of data with each data point representing the number of activated cells in an image. multiple images per mouse. Scale bars, 50 μm. Data are presented as means ± SEM. Statistical analysis was conducted using an unpaired two-tailed Mann Whitney test, with significance levels indicated as follows: ****P<0.0001.

**SI Figure 10.**
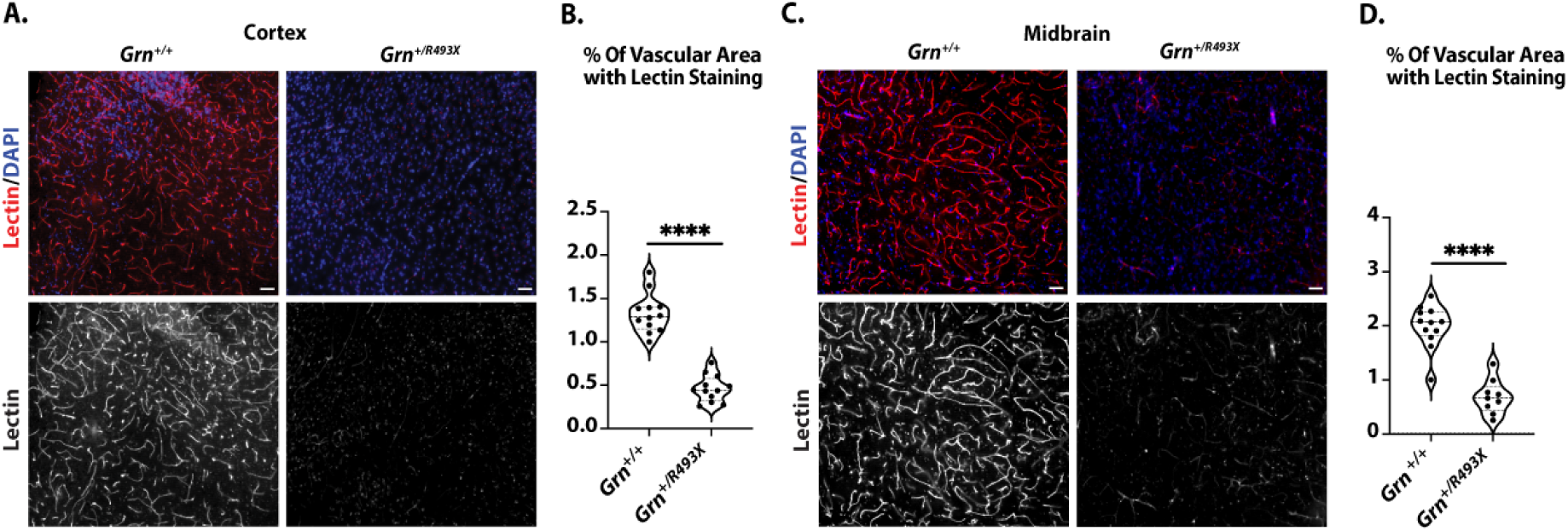
Evaluation of Tomato-Lectin Perfusion. **(A)**The representative immunofluorescence images of Tomato-lectin perfusion in mouse midbrain sections from 25–27-month old mice (n=3*Grn^R493X/+^* and littermate control mice (n=3)are shown. (B) Quantification of data, with each data point representing the fluorescence image intensity in one image. Multiple images per mouse. Scale bars, 50 μm. Data are presented as means ± SEM. Statistical analysis was conducted using an unpaired Mann Whitney test, with significance levels indicated as follows: ****P<0.0001

**SI Figure 11.**
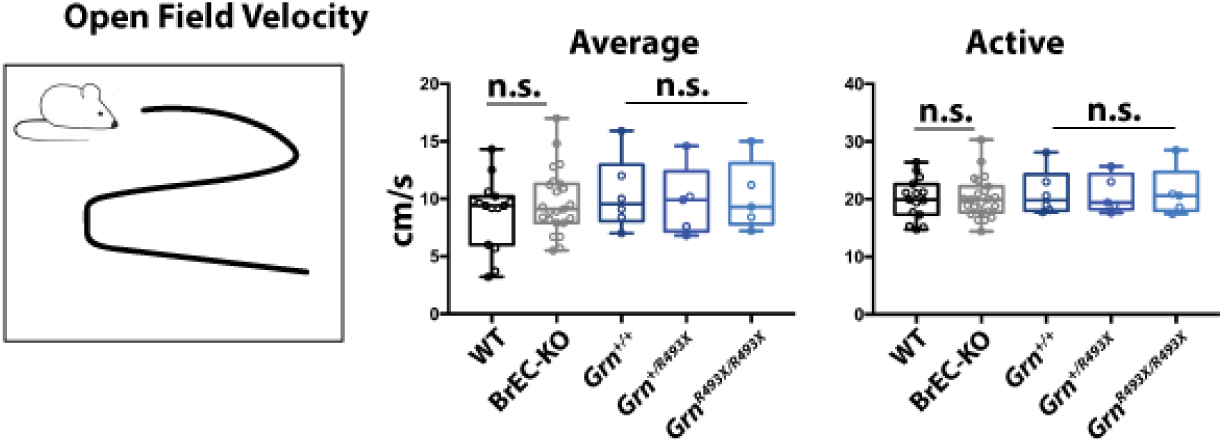
Behavioral Assessment of BrEC-KO Mice, Littermate Controls, and*Grn*+/R493X Knock-in Mouse Model of Frontotemporal Lobe Dementia (FTLD) in Open Field Velocity. BrEC-KO and littermate controls were assessed in behavioral core along with a knock-in mouse model of Frontal Temportal Lobe Dementia (FTLD, *Grn*+/R493X). No difference was observed in open field velocity.

## References

1. Tziortzouda P, Van Den Bosch L, Hirth F. Triad of TDP43 control in neurodegeneration: autoregulation, localization and aggregation. Nat Rev Neurosci. 2021 Apr;22(4):197–208.

2. Brown AL, Wilkins OG, Keuss MJ, Hill SE, Zanovello M, Lee WC, et al. TDP-43 loss and ALS-risk SNPs drive mis-splicing and depletion of UNC13A. Nature. 2022 Mar;603(7899):131–7.

3. Ma XR, Prudencio M, Koike Y, Vatsavayai SC, Kim G, Harbinski F, et al. TDP-43 represses cryptic exon inclusion in the FTD-ALS gene UNC13A. Nature. 2022 Mar;603(7899):124–30.

4. Ling JP, Pletnikova O, Troncoso JC, Wong PC. TDP-43 repression of nonconserved cryptic exons is compromised in ALS-FTD. Science. 2015 Aug 7;349(6248):650–5.

5. Melamed Z, López-Erauskin J, Baughn MW, Zhang O, Drenner K, Sun Y, et al. Premature polyadenylation-mediated loss of stathmin-2 is a hallmark of TDP-43-dependent neurodegeneration. Nat Neurosci. 2019 Feb;22(2):180–90.

6. Sabatelli M, Zollino M, Conte A, Del Grande A, Marangi G, Lucchini M, et al. Primary fibroblasts cultures reveal TDP-43 abnormalities in amyotrophic lateral sclerosis patients with and without SOD1 mutations. Neurobiol Aging. 2015 May;36(5):2005 e5–2005 e13.

7. Araki K, Araki A, Honda D, Izumoto T, Hashizume A, Hijki ata Y, et al. TDP-43 regulates early-phase insulin secretion via CaV1.2-mediated exocytosis in islets. J Clin Invest. 2019 Jul 29;130.

8. Serio A, Bilican B, Barmada SJ, Ando DM, Zhao C, Siller R, et al. Astrocyte pathology and the absence of non-cell autonomy in an induced pluripotent stem cell model of TDP-43 proteinopathy. Proc Natl Acad Sci U S A. 2013 Mar 19;110(12):4697–702.

9. Peng AYT, Agrawal I, Ho WY, Yen YC, Pinter AJ, Liu J, et al. Loss of TDP-43 in astrocytes leads to motor deficits by triggering A1-like reactive phenotype and triglial dysfunction. Proc Natl Acad Sci U S A. 2020 Nov 17;117(46):29101–12.

10. Licht-Murava A, Meadows SM, Palaguachi F, Song SC, Jackvony S, Bram Y, et al. Astrocytic TDP-43 dysregulation impairs memory by modulating antiviral pathways and interferon-inducible chemokines. Sci Adv. 2023 Apr 21;9(16):eade1282.

11. Montagne A, Barnes SR, Sweeney MD, Halliday MR, Sagare AP, Zhao Z, et al. Blood-brain barrier breakdown in the aging human hippocampus. Neuron. 2015 Jan 21;85(2):296–302.

12. Nation DA, Sweeney MD, Montagne A, Sagare AP, D’Orazio LM, Pachicano M, et al. Blood-brain barrier breakdown is an early biomarker of human cognitive dysfunction. Nat Med. 2019 Feb;25(2):270–6.

13. van de Haar HJ, Burgmans S, Jansen JF, van Osch MJ, van Buchem MA, Muller M, et al. Blood-Brain Barrier Leakage in Patients with Early Alzheimer Disease. Radiology. 2017 Feb;282(2):615.

14. Sagare AP, Bell RD, Zhao Z, Ma Q, Winkler EA, Ramanathan A, et al. Pericyte loss influences Alzheimer-like neurodegeneration in mice. Nat Commun. 2013;4:2932.

15. Mendiola AS, Yan Z, Dixit K, Johnson JR, Bouhaddou M, Meyer-Franke A, et al. Defining blood-induced microglia functions in neurodegeneration through multiomic profiling. Nat Immunol. 2023 Jul;24(7):1173–87.

16. Crouch EE, Doetsch F. FACS isolation of endothelial cells and pericytes from mouse brain microregions. Nat Protoc. 2018 Apr;13(4):738–51.

17. Ruck T, Bittner S, Epping L, Herrmann AM, Meuth SG. Isolation of primary murine brain microvascular endothelial cells. J Vis Exp. 2014 Nov 14;(93):e52204.

18. Heithoff BP, George KK, Phares AN, Zuidhoek IA, Munoz-Ballester C, Robel S. Astrocytes are necessary for blood-brain barrier maintenance in the adult mouse brain. Glia. 2021 Feb;69(2):436–72.

19. Sweeney MD, Sagare AP, Zlokovic BV. Blood-brain barrier breakdown in Alzheimer disease and other neurodegenerative disorders. Nat Rev Neurol. 2018 Mar;14(3):133–50.

20. Petersen MA, Ryu JK, Akassoglou K. Fibrinogen in neurological diseases: mechanisms, imaging and therapeutics. Nat Rev Neurosci. 2018 May;19(5):283–301.

21. Daneman R, Prat A. The blood-brain barrier. Cold Spring Harb Perspect Biol. 2015 Jan 5;7(1):a020412.

22. Claesson-Welsh L, Dejana E, McDonald DM. Permeability of the Endothelial Barrier: Identifying and Reconciling Controversies. Trends Mol Med. 2021 Apr;27(4):314–31.

23. Ridder DA, Lang MF, Salinin S, Roderer JP, Struss M, Maser-Gluth C, et al. TAK1 in brain endothelial cells mediates fever and lethargy. J Exp Med. 2011 Dec 19;208(13):2615–23.

24. Ayala YM, Misteli T, Baralle FE. TDP-43 regulates retinoblastoma protein phosphorylation through the repression of cyclin-dependent kinase 6 expression. Proc Natl Acad Sci U S A. 2008 Mar 11;105(10):3785–9.

25. Paul J, Strickland S, Melchor JP. Fibrin deposition accelerates neurovascular damage and neuroinflammation in mouse models of Alzheimer’s disease. J Exp Med. 2007 Aug 6;204(8):1999–2008.

26. Merlini M, Rafalski VA, Rios Coronado PE, Gill TM, Ellisman M, Muthukumar G, et al. Fibrinogen Induces Microglia-Mediated Spine Elimination and Cognitive Impairment in an Alzheimer’s Disease Model. Neuron. 2019 Mar 20;101(6):1099–1108 e6.

27. Nguyen AD, Nguyen TA, Zhang J, Devireddy S, Zhou P, Karydas AM, et al. Murine knockin model for progranulin-deficient frontotemporal dementia with nonsense-mediated mRNA decay. Proc Natl Acad Sci U A. 2018 Mar 20;115(12):E2849–58.

28. Brettschneider J, Toledo JB, Van Deerlin VM, Elman L, McCluskey L, Lee VMY, et al. Microglial activation correlates with disease progression and upper motor neuron clinical symptoms in amyotrophic lateral sclerosis. PloS One. 2012;7(6):e39216.

29. Krabbe G, Minami SS, Etchegaray JI, Taneja P, Djukic B, Davalos D, et al. Microglial NFkappaB-TNFalpha hyperactivation induces obsessive-compulsive behavior in mouse models of progranulin-deficient frontotemporal dementia. Proc Natl Acad Sci U A. 2017 May 9;114(19):5029–34.

30. Arrant AE, Filiano AJ, Warmus BA, Hall AM, Roberson ED. Progranulin haploinsufficiency causes biphasic social dominance abnormalities in the tube test. Genes Brain Behav. 2016 Jul;15(6):588–603.

31. Chou CC, Zhang Y, Umoh ME, Vaughan SW, Lorenzini I, Liu F, et al. TDP-43 pathology disrupts nuclear pore complexes and nucleocytoplasmic transport in ALS/FTD. Nat Neurosci. 2018 Feb;21(2):228–39.

32. Jeong YH, Ling JP, Lin SZ, Donde AN, Braunstein KE, Majounie E, et al. Tdp-43 cryptic exons are highly variable between cell types. Mol Neurodegener. 2017 Feb 2;12(1):13.

33. Schmid B, Hruscha A, Hogl S, Banzhaf-Strathmann J, Strecker K, van der Zee J, et al. Loss of ALS-associated TDP-43 in zebrafish causes muscle degeneration, vascular dysfunction, and reduced motor neuron axon outgrowth. Proc Natl Acad Sci U A. 2013 Mar 26;110(13):4986–91.

34. Hipke K, Pitter B, Hruscha A, van Bebber F, Modic M, Bansal V, et al. Loss of TDP-43 causes ectopic endothelial sprouting and migration defects through increased fibronectin, vcam 1 and integrin α4/β1. Front Cell Dev Biol. 2023;11:1169962.

35. Jackman K, Kahles T, Lane D, Garcia-Bonilla L, Abe T, Capone C, et al. Progranulin deficiency promotes post-ischemic blood-brain barrier disruption. J Neurosci. 2013 Dec 11;33(50):19579–89.

36. Zamudio F, Loon AR, Smeltzer S, Benyamine K, Navalpur Shanmugam NK, Stewart NJF, et al. TDP-43 mediated blood-brain barrier permeability and leukocyte infiltration promote neurodegeneration in a low-grade systemic inflammation mouse model. J Neuroinflammation. 2020 Sep 26;17(1):283.

37. Xu X, Zhang C, Jiang J, Xin M, Hao J. Effect of TDP43-CTFs35 on Brain Endothelial Cell Functions in Cerebral Ischemic Injury. Mol Neurobiol. 2022 Jul;59(7):4593–611.

38. Yanagida K, Liu CH, Faraco G, Galvani S, Smith HK, Burg N, et al. Size-selective opening of the blood-brain barrier by targeting endothelial sphingosine 1-phosphate receptor 1. Proc Natl Acad Sci U A. 2017 Apr 25;114(17):4531–6.

39. Nikolakopoulou AM, Montagne A, Kisler K, Dai Z, Wang Y, Huuskonen MT, et al. Pericyte loss leads to circulatory failure and pleiotrophin depletion causing neuron loss. Nat Neurosci. 2019 Jul;22(7):1089– 98.

40. Wood M, Quinet A, Lin YL, Davis AA, Pasero P, Ayala YM, et al. TDP-43 dysfunction results in R-loop accumulation and DNA replication defects. J Cell Sci. 2020 Oct 30;133(20):jcs244129.

41. Bodin A, Greibill L, Gouju J, Letournel F, Pozzi S, Julien JP, et al. Transactive response DNA-binding protein 43 is enriched at the centrosome in human cells. Brain J Neurol. 2023 Sep 1;146(9):3624–33.

42. Zhao W, Beers DR, Bell S, Wang J, Wen S, Baloh RH, et al. TDP-43 actvi ates microglia through NF-κB and NLRP3 inflammasome. Exp Neurol. 2015 Nov;273:24–35.

